# Master control of protein secretion by *Mycobacterium tuberculosis*

**DOI:** 10.1101/2025.03.13.643117

**Authors:** Rashmi Ravindran Nair, Virginia Meikle, Swati Dubey, Mikhail Pavlenok, Michael Niederweis

**Author notes:** Corresponding author: Michael Niederweis.

## Abstract

Tuberculosis is the leading cause of death from a single infectious disease. *Mycobacterium tuberculosis* secretes proteins using five ESX systems with distinctive functions essential for its growth and virulence. Here we show that a non-canonical supercomplex of the EsxU-EsxT proteins, encoded in the *esx-4* locus, with the orphan EsxE-EsxF proteins, encoded in the *cpnT* operon, is required for toxin secretion by *M. tuberculosis*. Surprisingly, the outer membrane localization of all Esx proteins and their secretion into the cytosol of infected macrophages also depend on the EsxEF-EsxUT supercomplex and ESX-4. These results not only demonstrate that the Esx proteins have dual functions as the long-sought outer membrane components of ESX systems and as secreted effector proteins, but also reveal a novel master control mechanism of protein secretion in *M. tuberculosis*. The mutual dependency of EsxEF and EsxUT on each other synchronizes ESX effector protein secretion, enabling *M. tuberculosis* to block phagosomal maturation and to permeabilize the phagosomal membrane only when it is capable of killing host cells by toxin secretion. The requirement of the ESX-4 system for general protein secretion is a critical vulnerability which could be targeted by drugs and/or vaccines to simultaneously block many virulence factors of *M. tuberculosis*.

## INTRODUCTION

An estimated 10.8 million people fell ill with tuberculosis worldwide in 2023^1^. This devastating disease is caused by *Mycobacterium tuberculosis* and is the world’s leading cause of death from a single infectious disease killing approximately 1.25 million people in 2023^1^. Treatment of tuberculosis requires a combination therapy with 4 drugs and lasts 6-9 months^1^. Multi-drug resistant (MDR) strains of *M. tuberculosis* cause ∼4% of new active TB cases and are even more challenging to treat^1^. Thus, there is a critical need to better understand the molecular mechanisms which enable *M. tuberculosis* to survive inside the human body and cause disease. Secretion of proteins is essential for bacteria to acquire nutrients, to adapt to different environments and, during an infection, to survive, subvert and kill host cells^2,3^. The only know protein secretion systems in *M.* are the five type VII or ESX secretion systems. ESX-1, ESX-2 and ESX-4 are required to permeabilize the phagosomal membrane, a critical step in the survival and escape of *M. tuberculosis* from macrophages^4^. This process is essential for virulence of *M. tuberculosis*^5^. The ESX-3 system is required by *M. tuberculosis* to acquire iron^6^ and zinc^7^ and to inhibit phagosomal membrane repair^8^, while ESX-5 is required for growth, probably by controlling PPE and other proteins involved in nutrient uptake^9–13^. The ESX-4 system is also required for secretion of TNT^4^, the only known exotoxin and main cytotoxicity factor of *M. tuberculosis* in macrophages^14^, into the cytosol of infected cells where it triggers necroptotic cell death^15,16^. These examples underline the critical role of the ESX systems for virulence and pathogenicity of *M. tuberculosis*.

The ESX substrate proteins belong to distinctive families classified as Esx, PE/PPE and Esp proteins^17^, which share either an T7SS signal sequence (YXXXD/E)^18^ and/or a WXG motif in Esx proteins^19^. Esx proteins are encoded in a pair of genes and are secreted as heterodimers forming a four-helix bundle^20^. Despite these similarities, each ESX system secretes a specific subset of substrate proteins exerting distinct physiological functions. The specificity of substrate proteins for each ESX system is established by cognate EspG chaperones for PE/PPE proteins^21^, and by a PE/PPE heterodimer associated with each ESX system for Esx proteins^22^. Significant progress towards a molecular understanding of mycobacterial protein secretion has also been made by solving the structures of the inner membrane core complex of the ESX-3^23^ and ESX-5 systems^24,25^. However, the outer membrane components enabling the secretion of ESX-substrate proteins remain unknown^17,20^.

In this study we examined how the ESX-4 system secretes the *M. tuberculosis* toxin. We show that the ESX-4 secretion substrates EsxU and EsxT form an outer membrane complex, which is not only required for toxin secretion, but also for secretion of all other Esx effector proteins. Surprisingly, we observed the same phenotype for the EsxE and EsxF proteins, which are encoded by orphan *esx* genes located in the toxin operon (Fig. 1a). We then show that EsxE-EsxF and EsxU-EsxT form a non-canonical protein complex in the outer membrane of *M. tuberculosis*, which is essential for the function of the ESX-4 system and for secretion of all Esx effector proteins. The discovery of a posttranslational master control mechanism of protein secretion by *M. tuberculosis* has profound implications for our understanding of *M. tuberculosis* pathogenicity and reveals an attractive target for the development of drugs and/or vaccines to simultaneously inactivate many virulence pathways of *M. tuberculosis*.

**Fig. 1:**
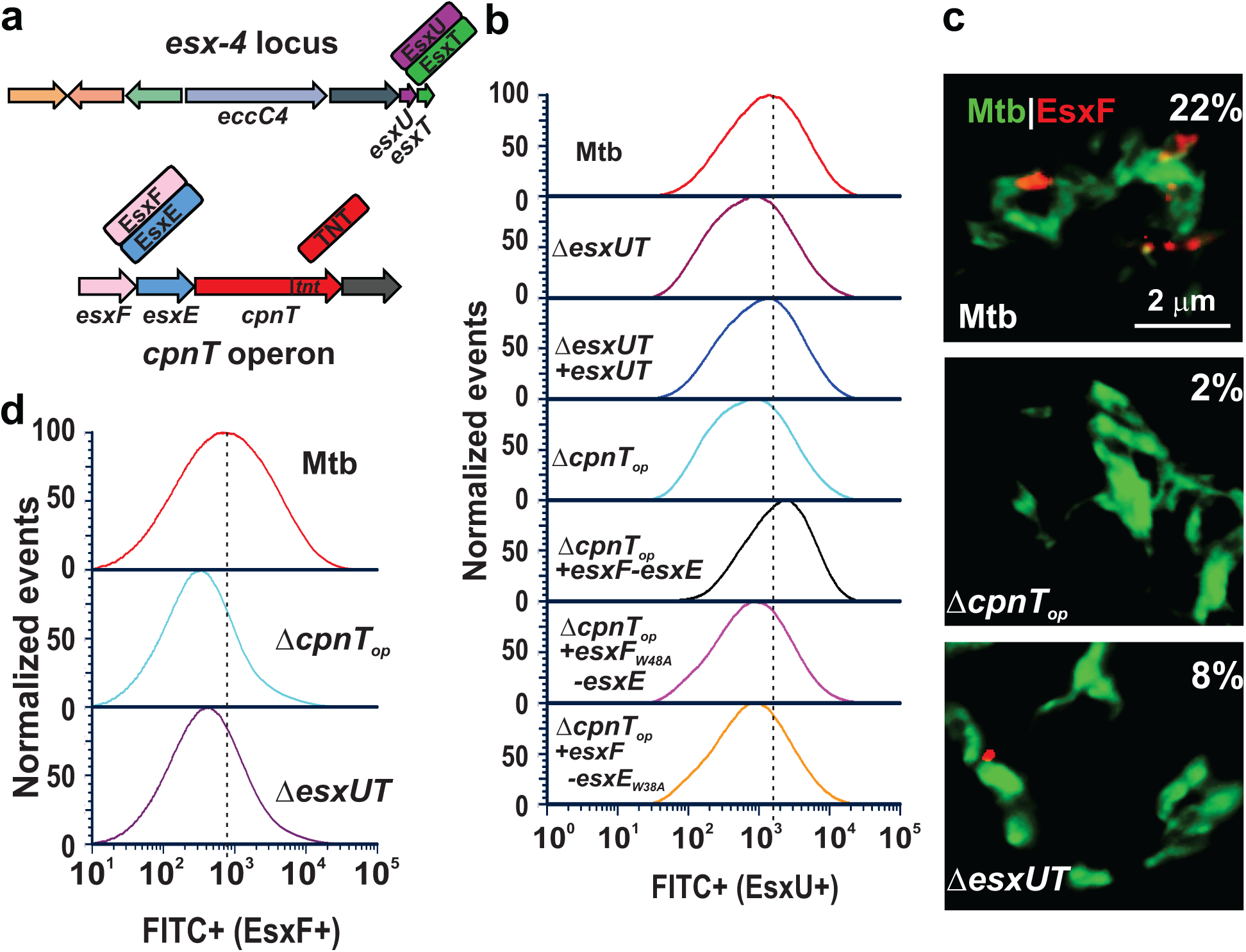
The surface accessibility of EsxEF and EsxUT in *M. tuberculosis* is mutually dependent on each other and requires pore formation by EsxEF. **a.** Genes located in the *esx-4* locus and the *cpnT* operon. The EsxE and EsxF proteins^28^ and the EsxU and EsxT proteins form heterodimers (see Fig. 2). **b.** Surface accessibility of EsxU in 50,000 *M. tuberculosis* cells by flow cytometry using an anti-EsxU antiserum. The peak fluorescence for wt *M. tuberculosis* is indicated by a dashed line. The strain names are indicated. The Δ*esxUT* mutant was complemented with the *esxUT* operon. The *cpnT* operon deletion mutant (Δ*cpnT*_op_) was partially complemented with the *esxFE* operon and mutated *esxFE* operons expressing the EsxF-W48A and EsxE-W38A mutants which are deficient in pore formation^28^. **c.** Detection of surface exposed EsxF in *M. tuberculosis* using fluorescence microscopy. The indicated *M. tuberculosis* strains were metabolically labeled with DMN-Trehalose (green) and stained with an anti-EsxF antiserum (red). The percentage of EsxF-positive cells relative to *M. tuberculosis* Δ*cpnT_op_* is indicated. **d.** Quantification of surface-accessible EsxF in 50,000 *M. tuberculosis* cells by flow cytometry using the EsxF antiserum.

## RESULTS

### EsxU is associated with membranes and is essential for toxin secretion by *M. tuberculosis*

Previously, we showed that the ESX-4 secretion system is essential for export of the CpnT channel protein, and for secretion its C-terminal toxin domain TNT by *M. tuberculosis*^4^. Since the molecular mechanism of outer membrane assembly is unknown for any mycobacterial protein, we aimed to identify the ESX-4 components required for this process and examined the EsxU and EsxT proteins, whose function in *M. tuberculosis* has not been described yet (Fig. 1a). To this end, we generated antisera against a synthetic EsxU peptide. Subcellular localization experiments reveal that EsxU is exclusively associated with membranes in cell lysates of wt *M. tuberculosis*, while only background levels are observed in *M. tuberculosis* strains lacking the *esxUT* genes or the gene the EccC4 ATPase which is essential for the activity of the ESX-4 system^4^ (Fig. S1a), demonstrating that EsxU is a substrate of the ESX-4 system. Considering its membrane association we hypothesized that EsxU in complex with its partner protein EsxT^26^ (Fig. 1a) are components of the ESX-4 system and might play a role in the ESX-4–mediated export of CpnT and TNT secretion. Fluorescence microscopy using protein-specific antibodies showed, as expected, that TNT is detectable on the cell surface of the parent *M. tuberculosis* strain in contrast to the deletion mutants lacking the *cpnT* operon (Δ*cpnT_op_*) and the *eccC4* gene (Fig. S1b, c). Surprisingly, surface-accessible TNT was also not detected in the *esxUT* deletion mutant (Fig. S1b, c). Quantitative analysis of ∼50,000 *M. tuberculosis* cells by flow cytometry using the TNT antiserum also showed a greatly reduced fluorescence for the Δ*cpnT_op_*, Δ*eccC4* and Δ*esxUT* mutants compared to the parent *M. tuberculosis* strain (Fig. S1c). Complementation with an *esxUT* expression plasmid restored the surface-accessibility of TNT in the *esxUT* deletion mutant to wt levels in the fluorescence microscopy (Figs. S1a, b) and flow cytometry experiments (Fig. S1c) demonstrating that EsxU and EsxT play an important role in TNT secretion.

Since *cpnT* expression in *M. tuberculosis* is strongly induced in macrophages compared to *in vitro* grown bacteria^15^, we determined TNT levels secreted into the cytosol of THP-1 cells infected with *M. tuberculosis*. Fluorescence microscopy revealed that wt *M. tuberculosis* secretes copious amounts of TNT into the cytosol of infected macrophages in contrast to the *esxUT* deletion mutant in which TNT was not detectable anymore similar to the *cpnT* operon deletion mutant. Complementation with *esxUT* fully restored TNT levels to wt levels (Figs. S1d, e). These experiments demonstrate that the surface accessibility and secretion of TNT into the cytosol of macrophages depends on the small ESX-4 substrate proteins EsxU and EsxT indicating an intriguing functional role in CpnT export and TNT secretion as components of the ESX-4 system, in contrast to the function of EsxUT as secreted effector proteins in *M. abscessus* ^27^.

### EsxU has a dual role as an outer membrane component of the ESX-4 system and as a secreted protein

Since EsxU is associated with membranes and is required for CpnT export and TNT secretion we hypothesized that EsxU is an essential component of the ESX-4 system. Indeed, fluorescence microscopy revealed that EsxU is surface accessible in *M. tuberculosis*, while only background fluorescence was observed in the Δ*eccC4* and Δ*esxUT* mutants (Fig. S2a, b). Quantitative analysis by flow cytometry confirmed the absence of EsxU in the *eccC4* and *esxUT* deletion mutants (Fig. S2c). In both experiments complementation of the Δ*esxUT* and the Δ*eccC4* mutants with integrative vectors expressing *esxUT* and *esx-4*, respectively, fully restored the surface accessibility of EsxU to wt levels (Fig. S2a, b). The combination of surface accessibility with membrane association shows that EsxU is anchored in the outer membrane as shown previously for other proteins of *M. tuberculosis*^10,11,14,28^. This conclusion is consistent with the essential function of EsxU in CpnT export and indicates that EsxU is an outer membrane component of the ESX-4 system.

This finding raised the question whether the function of EsxU is different from that of its homologous Esx proteins associated with the ESX systems of *M. tuberculosis*. For example, EsxB (CFP-10) is secreted by *M. tuberculosis* in a complex with the EsxA (ESAT-6) effector protein^29^, which plays an essential role in phagosomal permeabilization^5^. To examine whether EsxU is also a secreted effector protein, we infected THP-1 macrophages with *M. tuberculosis*. Fluorescence microscopy clearly shows both secreted EsxU and *M. tuberculosis* associated EsxU when macrophages are infected with wt *M. tuberculosis* but not with the *esxΔTU* and Δ*eccC4* strains (Fig. S2d, e). Wild-type levels of secreted EsxU were restored when the *esxUT* or *esx-4* genes were expressed in the respective mutants (Fig. S2d, e). Taken together, these results establish that EsxU is both a secreted effector protein and is an outer membrane component of the ESX-4 system required for CpnT export and TNT secretion.

### The surface localization of EsxU and EsxEF is co-dependent in *M. tuberculosis*

We previously reported that the EsxEF complex is essential for export of CpnT and secretion of TNT by *M. tuberculosis*^28^. Considering the surface localization of EsxUT in *M. tuberculosis* and its requirement for TNT export, we investigated whether EsxEF also contributes to the function of EsxUT. Consistent with this hypothesis, quantitative flow cytometry experiments show a substantial reduction of surface-accessible EsxU in the *cpnT* operon deletion mutant, similar to the level of the *esxUT* deletion mutant, compared to wt *M. tuberculosis* (Fig. 1b). Unexpectedly, fluorescence microscopy and flow cytometry experiments show that EsxU and EsxT are also essential for surface localization of EsxF (Fig. 1c, d). Taken together, these experiments demonstrate that both EsxEF and EsxUT are essential for CpnT export and TNT secretion and are co-dependent on each other to reach the cell surface of *M. tuberculosis* and to execute their functions.

### EsxUT oligomers do not appear to form membrane-spanning pores

We showed previously that EsxE and EsxF form an outer membrane pore, which is essential for CpnT export and toxin secretion by *M. tuberculosis*^28^. To investigate whether the homologous EsxUT proteins are also capable of forming pores, we purified the EsxUT complex based on our protocol developed for purification of EsxEF from *E. coli*^28^ (Fig. S3). While we observed the formation of oligomeric EsxUT proteins, extensive lipid bilayer experiments did not show significant channel activity by the EsxUT complex (Fig. S4 a-d). We then used microscopy to visualize fluorescent giant unilamellar vesicles, but did not observe any change of the vesicles in the presence of purified EsxUT (Fig. S4 e, f), in contrast to EsxEF which caused wide-spread vesicle destruction and deformation^28^. These experiments indicate that the EsxU-EsxT complex does not form membrane-spanning channels in contrast to EsxEF. However, considering the sensitivity of the channel activity of EsxEF to minor changes in buffer composition and pH^28^, we cannot exclude that EsxUT might be capable of pore formation under different experimental conditions.

### Pore formation by EsxEF is required for surface localization of EsxU

Previously, we observed that EsxE and EsxF proteins with WXG motif mutations (EsxF-W48A, EsxE-W38A) are membrane associated and accessible on the cell surface of *M. tuberculosis*, but are incapable of pore formation and do not translocate TNT to the cell surface^28^. To examine whether pore formation by EsxEF is required for the function of EsxU, we utilized the flow cytometry assay of *M. tuberculosis* cells. Surprisingly, no surface-accessible EsxU was detected in *M. tuberculosis* strains expressing *esxFE* operons encoding the EsxF-W48A and EsxE-W38A variants similar to the *M. tuberculosis esxUT* and *cpnT* operon deletion mutants (Fig. 1b). This key experiment demonstrates that pore formation by EsxEF is necessary for outer membrane association and surface accessibility of EsxU in *M. tuberculosis* establishing a central role for EsxEF in assembling a functional ESX-4 secretion system capable of translocating CpnT to the outer membrane and toxin secretion.

### The EsxEF and EsxUT proteins form a non-canonical complex in the outer membrane of *M. tuberculosis*, which is required for the function of the ESX-4 system

The results described above show that EsxEF and EsxU depend on each other for surface localization and for their function in TNT secretion. To investigate the hypothesis that EsxEF and EsxUT interact with each other, we co-expressed differentially tagged Esx proteins in the cytoplasm of *E. coli* using two compatible expression vectors encoding different pairs of Esx proteins (Fig. S5, Table S2). We chose *E. coli* as a host for these experiments to avoid the interference of functional ESX secretion systems in *M. tuberculosis* which would export and/or secrete Esx proteins. Pull-down assays were performed with *E. coli* lysates using histidine-tagged EsxF_His8_ as a bait protein captured on a nickel affinity column (Fig. 2a). Western blot analysis of the fraction eluted with imidazole not only detected EsxF and EsxE, as expected, but also EsxU and EsxT using tag-specific antibodies, indicating interactions between EsxE-EsxF and EsxU-EsxT (Fig. 2a). In a control experiment using EsxT-His_8_ as a bait protein we detected not only EsxU as expected, but also EsxE and EsxF (Fig. S6a). Thus, the results of both experiments consistently show that the EsxEF and EsxUT protein pairs form a complex in *E. coli*.

**Fig. 2.**
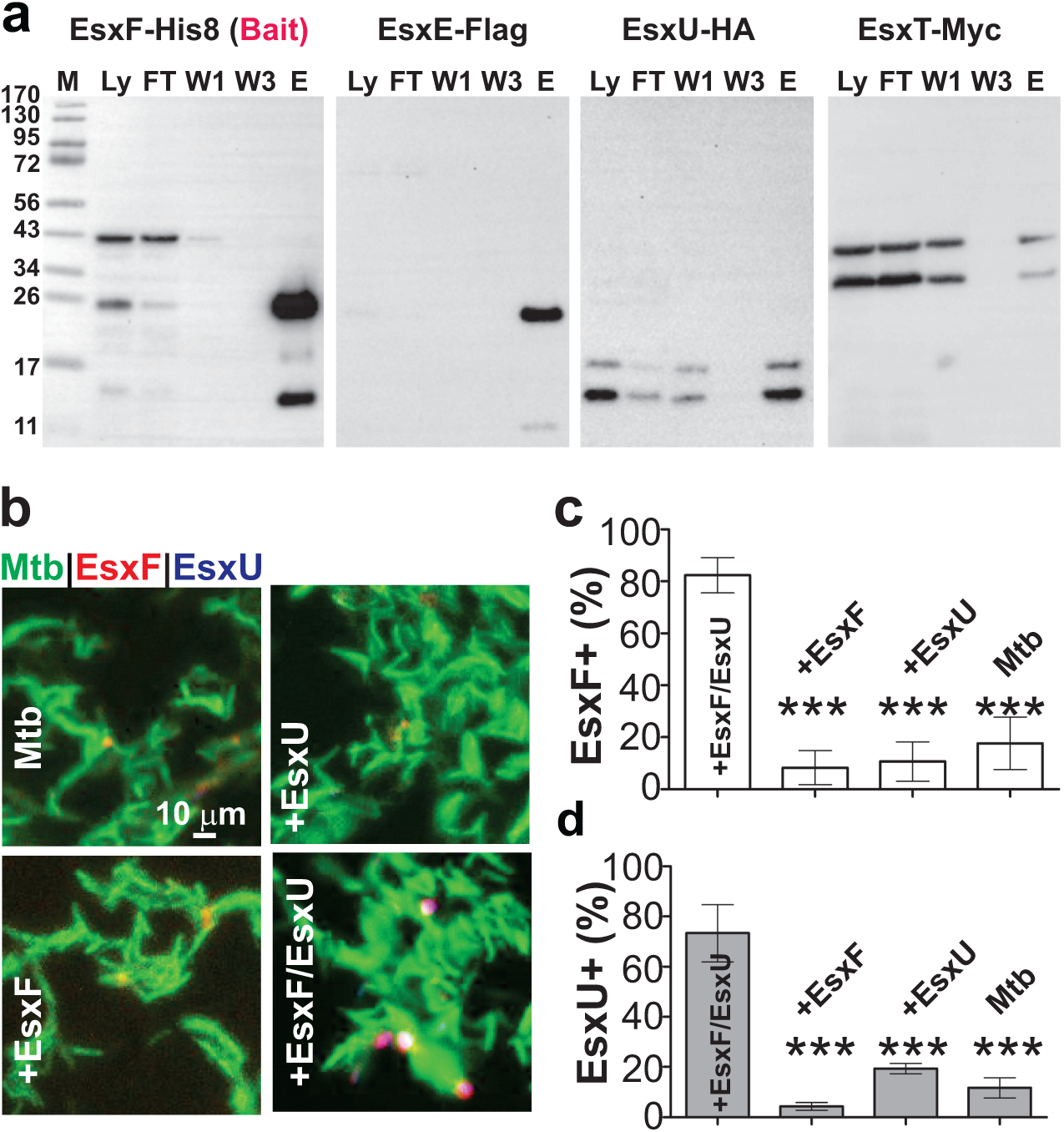
EsxEF and EsxUT interact and co-localize on the cell surface of *M. tuberculosis*. **a.** Pull-down assay: The bait protein EsxF_His8_ in a lysate (Ly) of *E. coli* producing EsxF_His8_/EsxE_FLAG_ and _HA_EsxU/EsxT_Myc_ was captured on a Ni(II) affinity column. The 10-fold diluted cell lysate (Ly), the flow through (FT), buffers after one wash (W1) and after three washes (W3) and the sample eluted with imidazol (E) were analyzed in Western blots using tag-specific antibodies. **b.** The surface accessibility of tagged EsxF and EsxU proteins in *M. tuberculosis* strains producing either _His8_EsxF-EsxE or _Myc_EsxU-EsxT or both Esx pairs together. Strains were scored positive for tagged EsxF or EsxU relative to untagged wt *M. tuberculosis*. The indicated *M. tuberculosis* strains were metabolically labeled with DMN-trehalose (green) and stained with anti-His (red) and anti-Myc (blue) Alexa fluor-conjugated antibodies. Co-localization of EsxF and EsxU in these *M. tuberculosis* strains was visualized by fluorescence microscopy showing the merged images and quantified as a percentage of positive cells (**c, d**). The single-channel fluorescence microscopy images are shown in Fig. S6.

Next, we examined whether EsxUT and EsxEF also interact with each other in their native environment, i.e. in the outer membrane of *M. tuberculosis*. To this end, we co-expressed operons encoding EsxE-EsxF_His8_ and EsxU_myc_-EsxT in *M. tuberculosis* (Table S1). Fluorescence microscopy showed that EsxF and EsxU indeed co-localize on the cell surface of *M. tuberculosis,* while no protein was detected in strains producing only one of these operons (Figs. 2b, c). Combined with the interaction experiments in *E. coli*, we conclude that EsxE-EsxF and EsxU-EsxT form a non-canonical Esx supercomplex in the outer membrane of *M. tuberculosis*. This complex formation explains the mutual requirement of these Esx proteins for their surface accessibility and for TNT secretion as shown in Fig. 1. Considering the requirement of the pore forming ability of EsxEF for formation of this complex (Fig. 1b), we hypothesize that the EsxEF-EsxUT complex forms the outer membrane pore which is required for the function of the ESX-4 secretion system.

### EsxEF and EsxUT are required for the surface exposure of other small Esx effector proteins

To investigate whether EsxEF and EsxUT play a more comprehensive role in export and/or secretion of small Esx proteins in *M. tuberculosis*, we expressed all *esx* genes associated with type VII secretion systems encoding HA-tagged proteins with their respective untagged partner proteins in the parent *M. tuberculosis* mc^2^6206 strain and in the Δ*cpnT_op_* (lacking *esxFE*) and Δ*esxUT* mutants. Fluorescence microscopy experiments revealed that HA-tagged EsxA and its homologs EsxC, EsxH, and EsxN are detected by HA antibodies on the cell surface of *M. tuberculosis* (Fig. 3), indicating that the Esx proteins encoded by the *esx-1*, esx-2, *esx-3* and *esx-5* loci, respectively, are also surface-accessible proteins and might be integral subunits of their respective ESX systems similar to the *esx-4* encoded EsxUT (Fig. 1). Surprisingly, the quantities of surface-accessible Esx proteins were greatly reduced in *M. tuberculosis* strains lacking the *cpnT* or the *esxUT* operons almost to the background level of the parent *M. tuberculosis* strain (Figs. 3, S7). Considering these intriguing results and the fact that the Esx proteins form heterodimers, we also examined the surface accessibility of their cognate partner proteins. Fluorescence microscopy shows that HA-tagged EsxB, EsxD, EsxG and EsxM are also surface-accessible in *M. tuberculosis* and their surface accessibility was dependent on EsxEF and EsxUT, albeit to a lesser extent than EsxA and its homologs (Figs. S8, S9). An important control experiment using an *M. tuberculosis* strain lacking *eccCa1-eccCb1* revealed that the surface exposure of EsxU was not compromised in the absence of a functional ESX-1 system in contrast to EsxA and EsxB secretion^29^, indicating a hierarchy of protein secretion in *M. tuberculosis* and an essential role of the ESX-4 system controlling protein secretion of the other ESX systems (Fig. S10). Taken together, these surprising results show that the surface accessibility of all small Esx proteins associated with the five ESX systems of *M. tuberculosis* depends to a large extent on the EsxEF-EsxUT supercomplex as an essential outer membrane component of the central ESX-4 secretion system of *M. tuberculosis*.

**Fig. 3:**
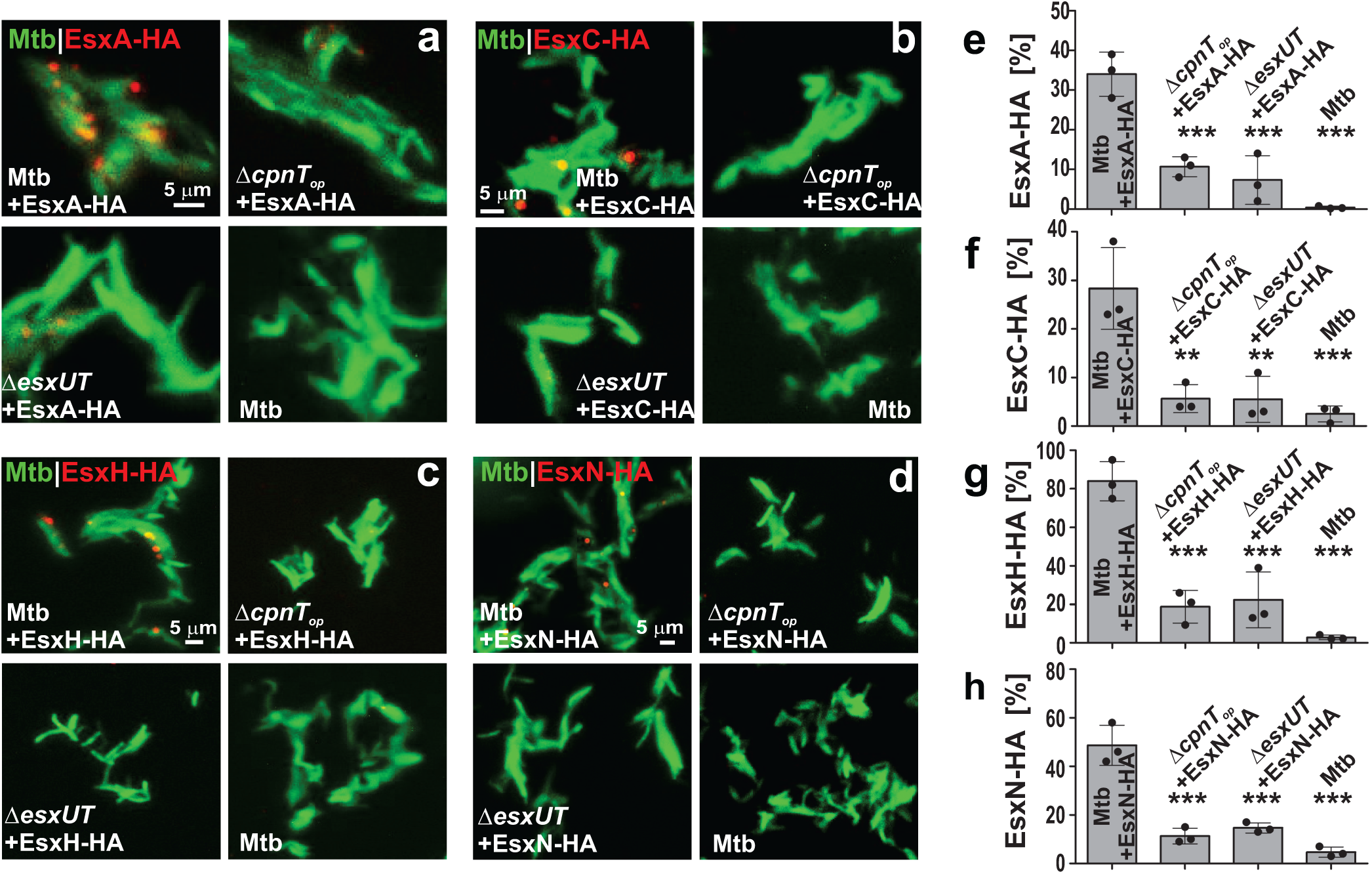
EsxEF and EsxUT are required for surface exposure of all Esx proteins of *M. tuberculosis*. Detection of surface-accessible Esx proteins of the different ESX systems in *M. tuberculosis* using fluorescence microscopy. The *M. tuberculosis cpnT* operon deletion (Δ*cpnT_op_*) mutant lacks the *esxF-esxE* genes. Scale bar = 5 µm. The *M. tuberculosis* strains produce different C-terminally HA-tagged Esx proteins as indicated: **(a)** EsxA (ESX-1), **(b)** EsxC (ESX-2), **(c)** EsxH (ESX-3) and **(d)** EsxN (ESX-5). The indicated *M. tuberculosis* strains were metabolically labeled with DMN-Trehalose (green), stained with a monoclonal HA-tag antibody (red) and were analyzed by fluorescent microscopy. The right column shows the quantification of surface-accessible Esx proteins in *M. tuberculosis* of the fluorescence images (n=3). Strains were scored positive for the respective Esx protein relative to wt *M. tuberculosis*, which does not produce an HA-tagged protein. Data are represented as mean ± SD. Asterisks indicate significant differences (** p ≤ 0.01, *** p ≤ 0.001; one-way ANOVA with Dunnett’s correction) in comparison to wt *M. tuberculosis* producing the same HA-tagged Esx protein.

### The EsxEF-EsxUT supercomplex controls effector protein secretion of other ESX systems in macrophages infected with *M. tuberculosis*

The paradigm-changing finding that the outer membrane EsxEF-EsxUT supercomplex is essential for protein secretion by all other ESX systems contradicts the current model that the effector proteins are solely dependent on their associated ESX systems^20,30^. Considering that *M. tuberculosis* is an obligate human pathogen and that the *esx-4* locus and the *cpnT* operon are strongly induced during infection^31^, we examined whether the central role of EsxEF-EsxUT in protein secretion by *M. tuberculosis* is also evident *in vivo*. To this end, we tracked the secretion of EsxA, EsxC and EsxU as representatives of their associated ESX-1, ESX-2 and ESX-4 systems, respectively, in infected macrophages. As expected and consistent with the *in vitro* results, EsxU secretion by *M. tuberculosis* in macrophages is dependent on EsxEF and vice versa (Fig. 4). Importantly, only background levels of EsxA (Fig. 4e, f) and EsxC (Fig. 4g, h) were secreted in strains lacking either EsxEF or EsxUT. Secretion was restored to wt levels upon complementation with *esxFE* or *esxUT* expression vectors. These results demonstrate that the EsxEF-EsxUT supercomplex is essential for protein secretion by other ESX systems of *M. tuberculosis* also during infection, consistent with the *in vitro* experiments. We conclude that the EsxEF-EsxUT supercomplex is essential for surface localization and secretion of small Esx proteins and possibly other proteins of the type VII secretion systems in *M. tuberculosis*.

**Fig. 4:**
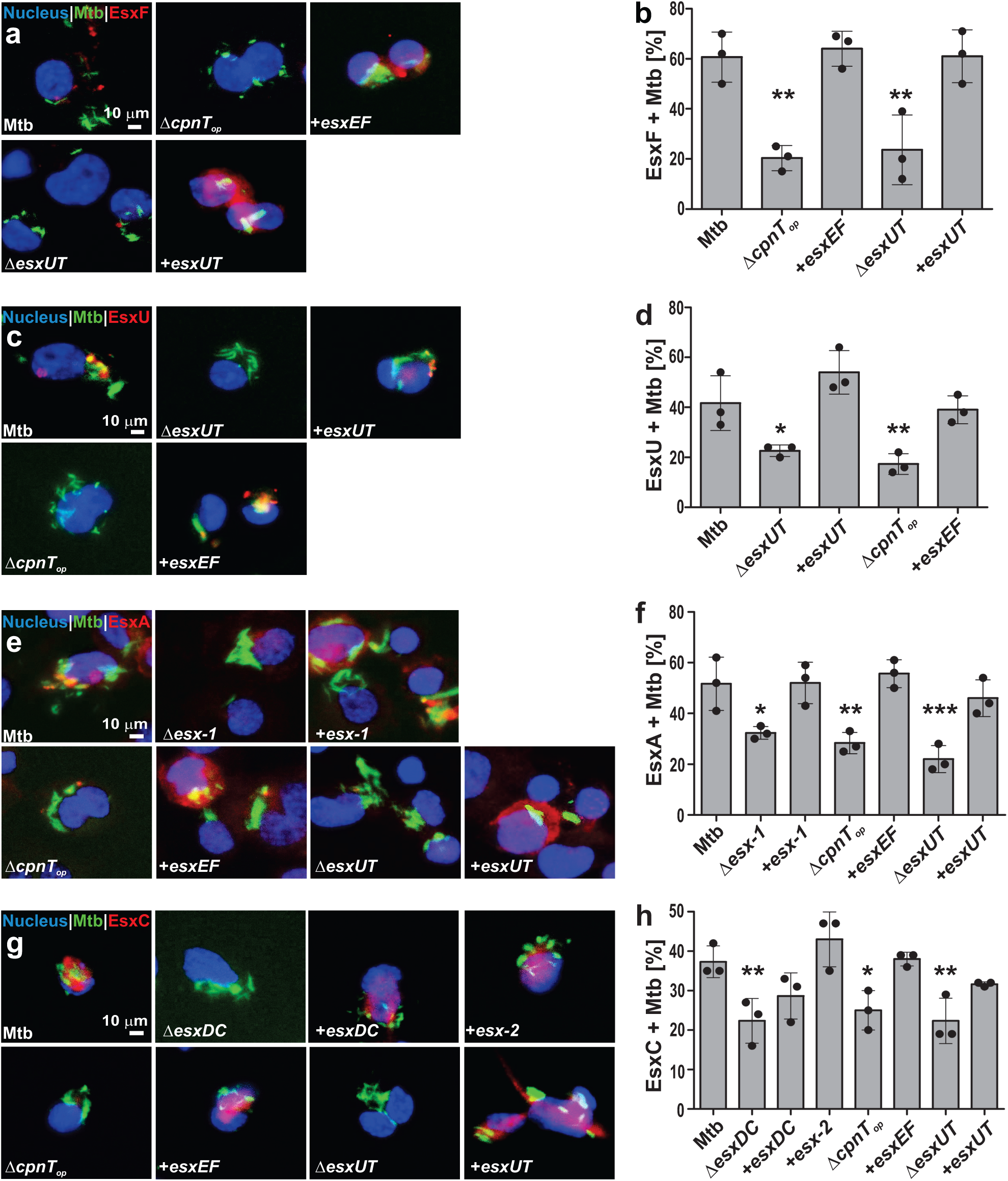
EsxEF and EsxUT are required for secretion of EsxA and EsxC into the cytosol of macrophages infected with *M. tuberculosis*. Analysis of secretion of Esx proteins into the cytosol of THP-1 macrophages infected with DMN-trehalose labelled *M. tuberculosis* strains (green) at a multiplicity of infection (MOI) of 10:1. The cells were fixed, permeabilized with 0.2% Triton X-100 and stained using antisera specific for EsxF (**a, b**), EsxU **(c, d**), EsxA (**e, f**) and EsxC (**g, h**). The nuclei of the macrophages were stained with DAPI (blue). Scale bar = 10 µm. The quantification of secretion of the respective Esx proteins into the cytosol of infected macrophages from the images shown in the left column (n=3). All quantitative data is represented as mean ± standard deviation (SD). Asterisks indicate significant differences (* p ≤ 0.05, ** p≤ 0.01, *** p ≤ 0.001; one-way ANOVA with Dunnett’s correction) in comparison to wt *M. tuberculosis*.

### EsxUT, EsxCD and EsxEF are secreted effector proteins of *M. tuberculosis* required for phagosomal permeabilization

We previously showed that the ESX-2 and ESX-4 systems are required for phagosomal permeabilization by *M. tuberculosis*^4^, in addition to the ESX-1 system^5,32,33^. Our infection experiments clearly show that EsxC and EsxU are secreted into the cytosol of macrophages (Fig. 4). However, it is unclear whether they have direct functions in phagosomal permeabilization similar to the ESX-1 effector proteins EsxA and EsxB^34,35^. To investigate the hypothesis, we constructed an *M. tuberculosis esxDC* deletion mutant (ML3000, Table S1) and infected THP-1 macrophages with parent *M. tuberculosis* and the respective mutants as indicated in Fig. 5 and determined the damage to the phagosomal membrane by these *M. tuberculosis* strains using Galectin-3 as a marker of ruptured host membranes^4,8,36^. We also determined the Galectin-3 levels in the *cpnT_op_* deletion mutant considering the important role of EsxEF in controlling the secretion of all other type VII effector proteins by *M. tuberculosis*. As expected, Galectin-3 staining shows substantial phagosomal membrane damage in THP-1 cells infected with the parent *M. tuberculosis* mc^2^6206 strain (Fig. 5). As a control we used the *M. tuberculosis eccCa1-eccCb1* deletion mutant (labeled Δ*esx-1*), which is impaired in EsxA and EsxB secretion^29^ and shows drastically reduced membrane damage (Fig. 5) consistent with previous publications^4,5,32^. The Galectin-3 levels of THP-1 cells infected with *M. tuberculosis* strains lacking *esxDC*, *esxUT* and *esxEF* displayed markedly reduced Galectin-3 levels similar to the level of the mutant with a defective ESX-1 system (Fig. 5). Importantly, the capability of these *M. tuberculosis* mutants to rupture phagosomal membranes was restored to wt levels by complementation with an ESX-1 cosmid or integrative vectors expressing *esxDC*, *esxUT* and *esxEF*. Similar results were obtained by determining the accessibility of antibodies in an antiserum against the tuberculin purified protein derivative (PPD) to the phagosome (Fig. S11). In conclusion, these results demonstrate that EsxCD, EsxUT and EsxEF are indeed essential for phagosomal permeabilization. However, these experiments do not reveal whether the role of EsxEF and EsxUT is indirect by controlling secretion of the other Esx effector proteins or whether the secreted EsxEF and EsxUT proteins directly participate in permeabilization of the phagosomal membrane.

**Fig. 5:**
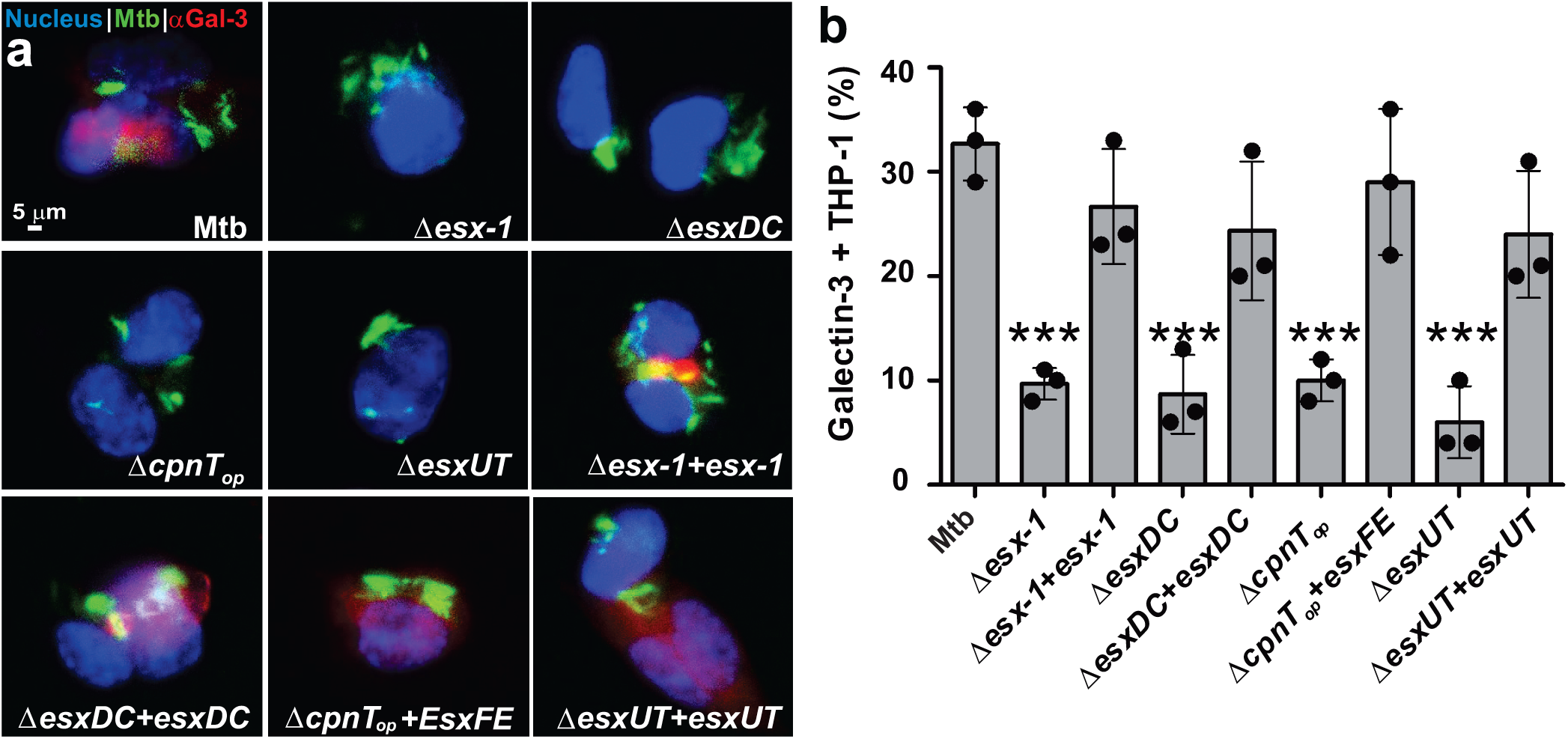
EsxEF, EsxUT and EsxCD are required for phagosomal permeabilization in macrophages infected with *M. tuberculosis*. **a**. Analysis of phagosomal permeabilization in THP-1 macrophages infected with the indicated *M. tuberculosis* strains at a multiplicity of infection (MOI) of 10:1. *M. tuberculosis* was metabolically labeled using DMN-Trehalose (green). The infected THP-1 cells were fixed, permeabilized with Triton X-100 and incubated with an Galectin-3 antibody (red) indicating membrane damage and with DAPI to visualize nuclei (blue). Scale bar = 5 µm. **b**. Quantification of infected macrophages positive for Galectin-3 compared to the *eccCa1-eccCb1* deletion mutant (labeled Δ*esx-1*) (n=3). Data are represented as mean ± SD. Asterisks indicate significant differences (*** p ≤ 0.001 calculated using the one-way ANOVA with Dunnett’s correction) in comparison to the wt strain.

## DISCUSSION

### The Esx proteins of *M. tuberculosis* share dual functions as outer membrane subunits of the ESX systems and as secreted effector proteins

In this study, we show that EsxU and EsxT, the small WXG100 proteins encoded in the *esx-4* locus (Fig. 6), form a complex which exists in two populations. One population is anchored in the outer membrane of *M. tuberculosis* and is essential for CpnT assembly in the outer membrane and TNT secretion, while another population is secreted into the cytosol of infected macrophages and is required for phagosomal permeabilization. Surface exposure and secretion of EsxU and EsxT are dependent on the ESX-4 system in contrast to *M. marinum*, which does not appear to secrete EsxU and EsxT^37^. A comprehensive analysis revealed that all Esx proteins associated with type VII secretion systems are surface exposed in *M. tuberculosis*. These results provide the first direct evidence of Esx proteins as outer membrane components of their respective ESX systems and explain recent proteomic studies revealing a requirement of EsxA for secretion of the ESX-1 substrates in *M. tuberculosis*^38^ and a hierarchical secretion of ESX-1 substrates depending on EsxA and EsxB in *M. marinum*^39,40^. A localization-specific proteomic study detected Esx proteins in the periplasm of *M. tuberculosis* consistent with their roles as outer membrane components of the ESX systems, although temporary exposure of secretion substrates could not be ruled out^41^. The location of the Esx proteins in the outer membrane and the capacity of some of these proteins to form membrane-spanning channels^28^ indicate that they are the long-sought channel proteins required for substrate translocation by the ESX systems across the outer membrane. It is possible that other proteins contribute to outer membrane translocation of ESX secretion substrates. For example, several PPE proteins have been identified as outer membrane proteins in *M. tuberculosis*^10,11^ and some have channel activity^42^ and/or transport functions^13^. However, no PPE proteins are encoded in the *esx-4* locus indicating that PPE proteins are not essential for ESX secretion. However, it is possible that the PPE proteins encoded in other *esx* loci and/or unknown proteins contribute to the function and/or efficiency of the ESX secretion machinery.

**Fig. 6:**
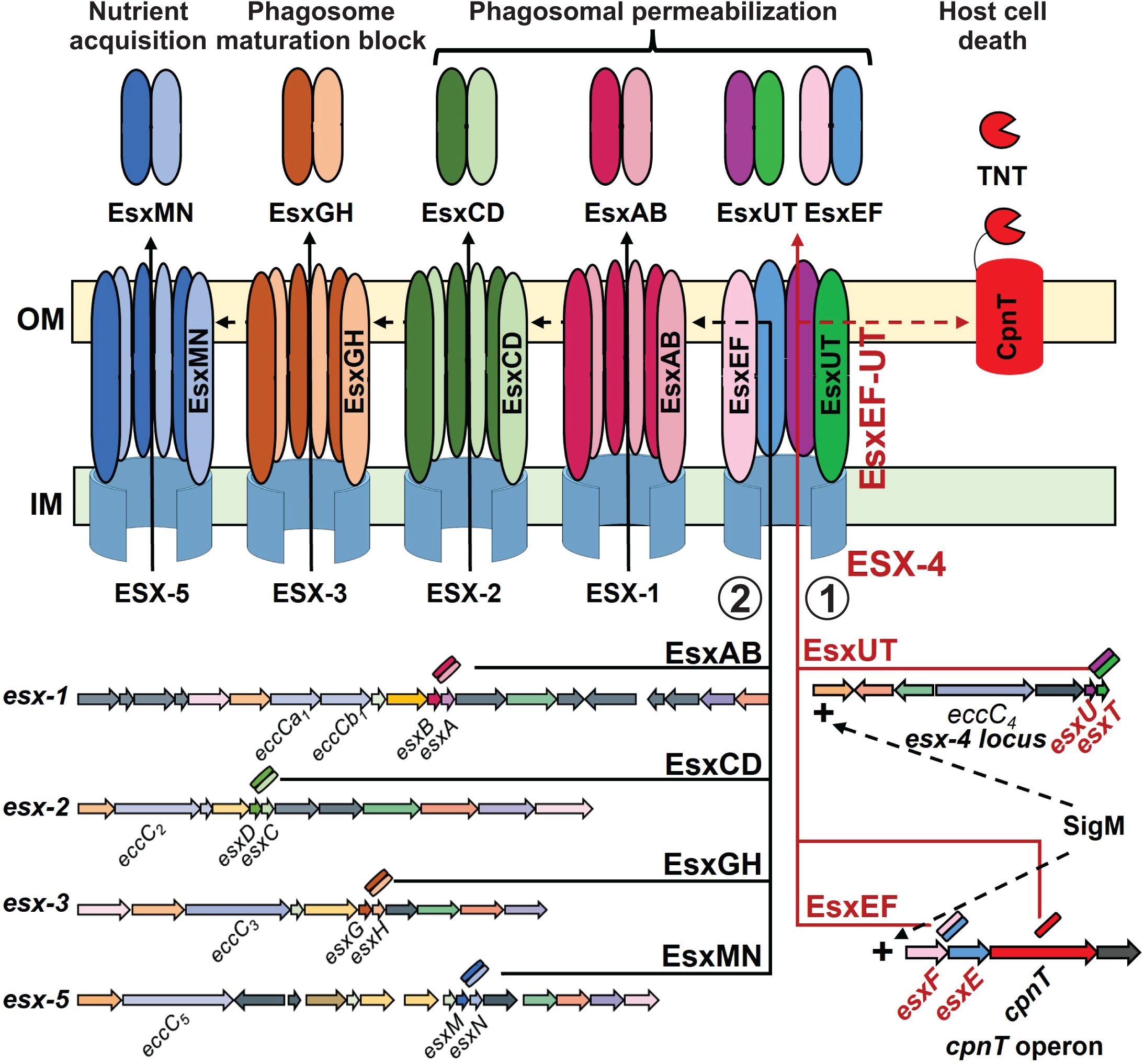
Unifying model of protein secretion in *M. tuberculosis*. SigM activates the expression of *esx4* locus and of the *cpnT* operon^64^. The ESX-4 core complex assembles in the inner membrane (IM) and exports the EsxEF-EsxUT complex to the *M. tuberculosis* cell surface. EsxUT forms a non-canonical complex with EsxEF. Pore formation by EsxEF is essential for the activity of the EsxEF-EsxUT supercomplex in the outer membrane in exporting CpnT and the small Esx proteins encoded by the *esx-1*, *esx-2*, *esx-3* and *esx-5* loci to the outer membrane. These small Esx proteins are detectable on the cell surface and may function as channels in the outer membrane facilitating the secretion of substrate proteins of each ESX system. EsxGH impairs phagosome maturation^43,69^. EsxAB, EsxCD, EsxUT and EsxEF permeabilize the phagosomal membrane. These processes enable TNT trafficking to the macrophage cytosol, where it hydrolyses NAD(P)^+^ and triggers necroptotic cell death of the infected macrophages^4^. Then, the substrate proteins of the ESX-5 system enable nutrient acquisition from the dying macrophage^11,70^.

Macrophage infection experiments show that the substrates of the ESX-1 (EsxA, EsxB), ESX-2 (EsxC, EsxD) and ESX-4 systems (EsxU, EsxT) are secreted. These results together with publications demonstrating secretion of the *esx-3* encoded EsxH and EsxG^43,44^ and *esx-5* encoded EsxN^9^ unequivocally show that all Esx proteins associated with the five type VII secretion systems of *M. tuberculosis* have dual functions as outer membrane subunits of their cognate ESX systems and as effector proteins. Since the ESX systems are the only known protein secretion systems in mycobacteria, these findings have profound implications for our molecular understanding of type VII protein secretion in the *Mycobacterium* genus, which contains over 190 species ^45^.

### EsxEF and EsxUT form an outer membrane supercomplex which is essential for toxin secretion in *M. tuberculosis*

An Esx protein sequence alignment shows that EsxU is the only Esx protein of *M. tuberculosis* lacking the highly conserved YxxxD/E secretion signal (Fig. S12), which is required for type VII secretion in mycobacteria^18,28,29^. However, our results clearly show that EsxU is translocated to the cell surface of *M. tuberculosis* and secreted into the cytosol of infected macrophages. This apparent contradiction is resolved by our finding that EsxUT and EsxEF form a non-canonical supercomplex and co-localize on the cell surface of *M. tuberculosis*. Thus, it appears that EsxUT has lost its capacity to be secreted, to interact with membranes and to form pores, but has gained the ability to interact with EsxEF. This potential is illustrated by high stability scores of Alphafold models of the canonical EsxEF and the heterologous EsxEU complex (Fig. S13, Table S5), which we can also detect in *E. coli* (Fig. 2). A key result of this study is that pore formation by EsxEF is essential for the function of the EsxEF-EsxUT super-complex in TNT secretion. Since EsxF has the critical YxxxD/E signal^28^ and the EsxEF complex interacts with membranes and forms pores^28^, and since pore formation is critical for the function and secretion of EsxUT, we hypothesize that EsxEF and EsxUT bind to each other before being exported by the ESX-4 system as a supercomplex and together form a pore in the outer membrane of *M. tuberculosis* enabling substrate secretion by ESX-4. In this model EsxUT contributes essential interactions with the EccC ATPase of the ESX-4 system similar to that its homolog EsxB with the ESX-1 EccC protein^46^, explaining the mutual dependency of EsxEF and EsxUT on each other. These results provide the first evidence of a non-canonical complex formation between two Esx heterodimers encoded by different genetic loci. Previously, complex formation was only observed between the tandem pairs of Esx proteins encoded in the same *esx* locus^26,47–50^.

### The outer membrane EsxEF-EsxUT supercomplex and ESX-4 control protein secretion in *M. tuberculosis*

The most surprising result of this study is that the outer membrane localization and the secretion of all Esx proteins associated with the type VII secretion systems of *M. tuberculosis* depend on the ESX-4 system and its outer membrane EsxEF-EsxUT supercomplex. While the ESX-4 system is the simplest type VII secretion system in *M. tuberculosis* and is considered the precursor of the other four ESX systems^51^, knowledge of its function in *M. tuberculosis* was limited to its requirement for phagosomal permeabilization^4^, toxin secretion^4^ and heme uptake^42^. In *M. abscessus* the ESX-4 system also mediates phagosomal rupture^52^, while it is required for conjugation in *M. smegmatis*^53^. By contrast, the ESX-4 system of *M. marinum* is considered to be non-functional^37^. These divergent functions or possible loss of functions of the ESX-4 systems appear to be the consequence of evolutionary adaptation by different mycobacteria to their particular ecological niches depending on the presence of other ESX systems after gene duplication of the ancient *esx-4* locus^54^. The low expression levels of the *esx-4* locus and the *cpnT* operon during *in vitro* growth of *M. tuberculosis* and the dependency of all type VII secretion on EsxEF-EsxUT raise the question how *M. tuberculosis* can sustain *in vitro* growth which requires the ESX-3 and ESX-5 systems for uptake of iron/zinc^7,55,56^ and other nutrients^57^, respectively. One possible explanation is that the low expression levels of the *esx-4* and *cpnT* operons are sufficient for basal activity of the other ESX systems. This conclusion is consistent with the observation of ESX protein secretion by *M. tuberculosis* grown *in vitro* as shown by the presence of EsxAB and EsxGH in culture supernatants^6,58,59^.

### Unifying model of protein secretion by *M. tuberculosis*

From this study and previous publications a unifying model has emerged. In the initial stages of the infection *M. tuberculosis* grows and replicates inside the lung. Then, CD4 T cells activate infected macrophages as part of the adaptive immune response including formation of granulomas to wall off the infection^60,61^. Under these adverse conditions the very low permeability of its outer membrane is crucial for survival of *M. tuberculosis*^62,63^. When growth of *M. tuberculosis* reaches a stationary phase, the alternative sigma factor SigM is induced which in turn activates expression of the *cpnT* operon and the *esx-4* locus (Fig. 6). This transcriptional control is highlighted by the observation that overexpression of *sigM* drastically induces transcription of the *esxFE* and *esxT* genes by 60-90-fold and of *esxU* by 1800-fold. SigM also induces its own transcription by 80-fold^64^. The simultaneous expression of *esxFE* and the *esx-4* operon including *esxUT* enables formation of the inner membrane ESX-4 machinery and the outer membrane EsxEF-EsxUT supercomplex and, subsequently, the export and secretion of other Esx substrates (Fig. 6). The delivery of the other outer membrane Esx proteins activates their respective ESX systems. Secretion of EsxGH by the ESX-3 system impairs phagosomal maturation^43^ and prepares *M. tuberculosis* for iron and zinc uptake^7,55^. Secretion of EsxAB and EsxCD by the ESX-1 and ESX-2 systems, respectively, together with EsxEF and EsxUT, ensures that toxin secretion is coordinated with the permeabilization of the phagosomal membrane. The rupture of the phagosomal membrane enables access of TNT and other secreted proteins such as PknG^65^ and SapM^66^ to the cytoplasm of the infected macrophages. TNT then degrades NAD^+15^, which in turn kills the macrophages by inducing necroptosis^16^. The following extensive membrane damage of the dying macrophage provides *M. tuberculosis* with the nutrients in the host cytoplasm. These nutrients are taken up by the PPE proteins^10,11,13^, which are exported to the outer membrane by the ESX-5 system^9,67^. The repression of the biosynthesis of the outer membrane lipid phthiocerol dimycocerosate (PDIM) by SigM and the following outer membrane remodeling by inducing the synthesis of other lipids^64^ is likely to benefit nutrient uptake through the PPE proteins and through the pore-forming N-terminal domain of CpnT^14^.

In conclusion, the paradigm-changing discoveries of the dual functions of the Esx proteins as secreted effector proteins and as essential outer membrane components of the ESX systems and the master control of protein secretion by *M. tuberculosis* through the ESX-4 system with its non-canonical EsxEF-EsxUT outer membrane supercomplex are fundamental for understanding the molecular events enabling *M. tuberculosis* to survive and replicate in the human body and cause disease. The mutual dependency of EsxEF and EsxUT on each other synchronizes ESX effector protein secretion, enabling *M. tuberculosis* to block phagosomal maturation and to permeabilize the phagosomal membrane when it is ready to secrete the toxin and to kill host cells. The requirement of a single outer membrane protein complex for general protein secretion by *M. tuberculosis* is a critical vulnerability which could be targeted by drugs and/or vaccines to simultaneously block important virulence factors of *M. tuberculosis*. Such a strategy would avoid the low permeability of the outer membrane of *M. tuberculosis*, which is a key factor in its intrinsic drug resistance and a major hurdle in TB drug develepment^63,68^. Our new model of the coordinated protein secretion program by *M. tuberculosis* will certainly stimulate further exciting research in the tuberculosis field and, hopefully, new drug discovery efforts.

## MATERIAL AND METHODS

A detailed description of the material and methods used in this study is available in the supplement.

- Bacterial strains and growth conditions
- Plasmid construction
- Construction of gene deletion mutants of *M. tuberculosis*
- Phthiocerol dimycocerosate analysis in *M. tuberculosis*
- Generation of antibodies against small Esx proteins
- Detection of surface-accessible proteins in *M. tuberculosis* by flow cytometry
- Fluorescence microscope analysis of surface accessible proteins in *M. tuberculosis*
- THP-1 cell culture and differentiation
- Infection of macrophages with *M. tuberculosis*
- Fluorescence microscope analysis of macrophages infected with *M. tuberculosis*
- Preparation of *M. tuberculosis* cell lysate and subcellular localization
- Detection of proteins in Western blots
- Purification of the EsxU-EsxT protein complex
- Planar lipid bilayer experiments
- Membrane interaction assay using unilamellar vesicles
- Pull down assay using tagged small Esx proteins
- Co-localization analysis of Esx proteins in *M. tuberculosis*
- Statistical evaluation

## Supporting information

Complete Supplement

## ACKNOWLEDGMENTS

We thank Dr. Roland Brosch for providing the cosmid 2F9 and Dr. Avraneel Paul for help with the design of the model (Fig. 6) and for the Alphafold structure predictions. This work was supported by the National Institutes of Health grant R01 AI175106 to MN. The funding agency had no involvement in preparation of the article, study design, in the collection, analysis and interpretation of data, in the writing of the report and in the decision to submit the article for publication.

## AUTHOR CONTRIBUTIONS

MN and RRN conceived the study. RRN, VM, SD and MP designed and performed experiments. MN and RRN wrote the manuscript. All authors analyzed data and edited the manuscript.

## COMPETING INTERESTS

The authors declare no competing interests.

**This article contains supporting information.**

## REFERENCES

1 WHO. Global Tuberculosis Report. World Health Organization, Geneva, (2024).

2 Costa, T. R. et al. Secretion systems in Gram-negative bacteria: structural and mechanistic insights. Nat Rev Microbiol 13, 343–359, (2015).

3 Green, E. R. & Mecsas, J. Bacterial Secretion Systems: An Overview. Microbiol Spectr 4, VMBF-0012-2015, (2016).

4 Pajuelo, D. et al. Toxin secretion and trafficking by *Mycobacterium tuberculosis*. Nat Commun 12, 6592, (2021).

5 Simeone, R. et al. Phagosomal rupture by *Mycobacterium tuberculosis* results in toxicity and host cell death. PLoS Pathog 8, e1002507, (2012).

6 Tufariello, J. M. et al. Separable roles for *Mycobacterium tuberculosis* ESX-3 effectors in iron acquisition and virulence. Proc Natl Acad Sci U S A 113, E348–357, (2016).

7 Serafini, A., Pisu, D., Palu, G., Rodriguez, G. M. & Manganelli, R. The ESX-3 secretion system is necessary for iron and zinc homeostasis in *Mycobacterium tuberculosis*. PLoS One 8, e78351, (2013).

8 Mittal, E., et al. *Mycobacterium tuberculosis* Type VII Secretion System Effectors Differentially Impact the ESCRT Endomembrane Damage Response. MBio 9, e01765, (2018).

9 Bottai, D. et al. Disruption of the ESX-5 system of *Mycobacterium tuberculosis* causes loss of PPE protein secretion, reduction of cell wall integrity and strong attenuation. Mol Microbiol 83, 1195–1209, (2012).

10 Mitra, A., Speer, A., Lin, K., Ehrt, S. & Niederweis, M. PPE surface proteins are required for heme utilization by *Mycobacterium tuberculosis*. MBio 8, e01720, (2017).

11 Wang, Q. et al. PE/PPE proteins mediate nutrient transport across the outer membrane of *Mycobacterium tuberculosis*. Science 367, 1147–1151, (2020).

12 Boradia, V., Frando, A. & Grundner, C. The *Mycobacterium tuberculosis* PE15/PPE20 complex transports calcium across the outer membrane. PLoS Biol 20, e3001906, (2022).

13 Babu Sait, M. R., et al. PPE51 mediates uptake of trehalose across the mycomembrane of *Mycobacterium tuberculosis*. Sci Rep 12, 2097, (2022).

14 Danilchanka, O. et al. An outer membrane channel protein of *Mycobacterium tuberculosis* with exotoxin activity. Proc Natl Acad Sci U S A 111, 6750–6755, (2014).

15 Sun, J. et al. The tuberculosis necrotizing toxin kills macrophages by hydrolyzing NAD. Nat Struct Mol Biol 22, 672–678, (2015).

16 Pajuelo, D. et al. NAD(+) depletion triggers macrophage necroptosis, a cell death pathway exploited by *Mycobacterium tuberculosis*. Cell Rep 24, 429–440, (2018).

17 Bunduc, C. M., Bitter, W. & Houben, E. N. G. Structure and Function of the Mycobacterial Type VII Secretion Systems. Annu Rev Microbiol 74, 315–335, (2020).

18 Daleke, M. H. et al. General secretion signal for the mycobacterial type VII secretion pathway. Proc Natl Acad Sci U S A 109, 11342–11347, (2012).

19 Pallen, M. J. The ESAT-6/WXG100 superfamily--and a new Gram-positive secretion system? Trends Microbiol 10, 209–212, (2002).

20 Famelis, N., Geibel, S. & van Tol, D. Mycobacterial type VII secretion systems. Biol Chem 404, 691–702, (2023).

21 Daleke, M. H. et al. Specific chaperones for the type VII protein secretion pathway. J Biol Chem 287, 31939–31947, (2012).

22 Damen, M. P. M. et al. Modification of a PE/PPE substrate pair reroutes an Esx substrate pair from the mycobacterial ESX-1 type VII secretion system to the ESX-5 system. J Biol Chem 295, 5960–5969, (2020).

23 Famelis, N. et al. Architecture of the mycobacterial type VII secretion system. Nature 576, 321–325, (2019).

24 Beckham, K. S. H. et al. Structure of the mycobacterial ESX-5 type VII secretion system pore complex. Sci Adv 7, eabg9923, (2021).

25 Bunduc, C. M. et al. Structure and dynamics of a mycobacterial type VII secretion system. Nature 593, 445–448, (2021).

26 Pandey, H. et al. Biophysical and immunological characterization of the ESX-4 system ESAT-6 family proteins Rv3444c and Rv3445c from *Mycobacterium tuberculosis* H37Rv. Tuberculosis (Edinb) 109, 85–96, (2018).

27 Lagune, M. et al. The ESX-4 substrates, EsxU and EsxT, modulate *Mycobacterium abscessus* fitness. PLoS Pathog 18, e1010771, (2022).

28 Tak, U., Dokland, T. & Niederweis, M. Pore-forming Esx proteins mediate toxin secretion by *Mycobacterium tuberculosis*. Nat Commun 12, 394, (2021).

29 Champion, P. A., Stanley, S. A., Champion, M. M., Brown, E. J. & Cox, J. S. C-terminal signal sequence promotes virulence factor secretion in *Mycobacterium tuberculosis*. Science 313, 1632–1636, (2006).

30 Abdallah, A. M. et al. Type VII secretion--mycobacteria show the way. Nat Rev Microbiol 5, 883–891, (2007).

31 Hudock, T. A. et al. Hypoxia Sensing and Persistence Genes Are Expressed during the Intragranulomatous Survival of *Mycobacterium tuberculosis*. Am J Respir Cell Mol Biol 56, 637–647, (2017).

32 de Jonge, M. I. et al. ESAT-6 from *Mycobacterium tuberculosis* dissociates from its putative chaperone CFP-10 under acidic conditions and exhibits membrane-lysing activity. J Bacteriol 189, 6028–6034, (2007).

33 Tiwari, S., Casey, R., Goulding, C. W., Hingley-Wilson, S. & Jacobs, W. R., Jr. Infect and Inject: How *Mycobacterium tuberculosis* Exploits Its Major Virulence-Associated Type VII Secretion System, ESX-1. Microbiol Spectr 7, BAI-0024-2019, (2019).

34 Hsu, T. et al. The primary mechanism of attenuation of bacillus Calmette-Guerin is a loss of secreted lytic function required for invasion of lung interstitial tissue. Proc Natl Acad Sci USA 100, 12420–12425, (2003).

35 van der Wel, N. et al. *M.* *tuberculosis* and *M. leprae* translocate from the phagolysosome to the cytosol in myeloid cells. Cell 129, 1287–1298, (2007).

36 Thurston, T. L., Wandel, M. P., von Muhlinen, N., Foeglein, A. & Randow, F. Galectin 8 targets damaged vesicles for autophagy to defend cells against bacterial invasion. Nature 482, 414–418, (2012).

37 Wang, Y., et al. Crosstalk between the ancestral type VII secretion system ESX-4 and other T7SS in Mycobacterium marinum. iScience 25, 103585, (2022).

38 Toniolo, C., Dhar, N. & McKinney, J. D. Uptake-independent killing of macrophages by extracellular *Mycobacterium tuberculosis* aggregates. Embo J 42, e113490, (2023).

39 Champion, P. A., Champion, M. M., Manzanillo, P. & Cox, J. S. ESX-1 secreted virulence factors are recognized by multiple cytosolic AAA ATPases in pathogenic mycobacteria. Mol Microbiol 73, 950–962, (2009).

40 Cronin, R. M., Ferrell, M. J., Cahir, C. W., Champion, M. M. & Champion, P. A. Proteo-genetic analysis reveals clear hierarchy of ESX-1 secretion in *Mycobacterium marinum*. Proc Natl Acad Sci U S A 119, e2123100119, (2022).

41 Jaisinghani, N. et al. Proteomics from compartment-specific APEX2 labeling in *Mycobacterium tuberculosis* reveals Type VII secretion substrates in the cell wall. Cell Chem Biol 31, 523–533 e524, (2024).

42 Sankey, N., et al. Role of the *Mycobacterium tuberculosis* ESX-4 Secretion System in Heme Iron Utilization and Pore Formation by PPE Proteins. mSphere, e0057322, (2023).

43 Mehra, A., et al. *Mycobacterium tuberculosis* type VII secreted effector EsxH targets host ESCRT to impair trafficking. PLoS Pathog 9, e1003734, (2013).

44 Tinaztepe, E. et al. Role of Metal-Dependent Regulation of ESX-3 Secretion in Intracellular Survival of *Mycobacterium tuberculosis*. Infect Immun 84, 2255–2263, (2016).

45 Armstrong, D. T., Eisemann, E. & Parrish, N. A Brief Update on Mycobacterial Taxonomy, 2020 to 2022. J Clin Microbiol 61, e0033122, (2023).

46 Rosenberg, O. S. et al. Substrates Control Multimerization and Activation of the Multi-Domain ATPase Motor of Type VII Secretion. Cell 161, 501–512, (2015).

47 Renshaw, P. S. et al. Conclusive evidence that the major T-cell antigens of the *Mycobacterium tuberculosis* complex ESAT-6 and CFP-10 form a tight, 1:1 complex and characterization of the structural properties of ESAT-6, CFP-10, and the ESAT-6*CFP-10 complex. Implications for pathogenesis and virulence. J Biol Chem 277, 21598–21603, (2002).

48 Renshaw, P. S. et al. Sequence-specific assignment and secondary structure determination of the 195-residue complex formed by the Mycobacterium tuberculosis proteins CFP-10 and ESAT-6. J Biomol NMR 30, 225–226, (2004).

49 Renshaw, P. S. et al. Structure and function of the complex formed by the tuberculosis virulence factors CFP-10 and ESAT-6. Embo J 24, 2491–2498, (2005).

50 Ilghari, D. et al. Solution structure of the *Mycobacterium tuberculosis* EsxG.EsxH complex: functional implications and comparisons with other M. tuberculosis Esx family complexes. J Biol Chem 286, 29993–30002, (2011).

51 Roy, S., Ghatak, D., Das, P. & BoseDasgupta, S. ESX secretion system: The gatekeepers of mycobacterial survivability and pathogenesis. Eur J Microbiol Immunol (Bp) 10, 202–209, (2020).

52 Laencina, L. et al. Identification of genes required for *Mycobacterium abscessus* growth in vivo with a prominent role of the ESX-4 locus. Proc Natl Acad Sci U S A 115, E1002–E1011, (2018).

53 Gray, T. A. et al. Intercellular communication and conjugation are mediated by ESX secretion systems in mycobacteria. Science 354, 347–350, (2016).

54 Gray, T. A. & Derbyshire, K. M. Blending genomes: distributive conjugal transfer in mycobacteria, a sexier form of HGT. Mol Microbiol 108, 601–613, (2018).

55 Siegrist, M. S. et al. Mycobacterial Esx-3 is required for mycobactin-mediated iron acquisition. Proc Natl Acad Sci U S A 106, 18792–18797, (2009).

56 Serafini, A., Boldrin, F., Palu, G. & Manganelli, R. Characterization of a *Mycobacterium tuberculosis* ESX-3 conditional mutant: essentiality and rescue by iron and zinc. J Bacteriol 191, 6340–6344, (2009).

57 Elliott, S. R. & Tischler, A. D. Phosphate starvation: a novel signal that triggers ESX-5 secretion in *Mycobacterium tuberculosis*. Mol Microbiol 100, 510–526, (2016).

58 Bottai, D. et al. ESAT-6 secretion-independent impact of ESX-1 genes *espF* and *espG1* on virulence of *Mycobacterium tuberculosis*. J Infect Dis 203, 1155–1164, (2011).

59 Lim, Z. L., Drever, K., Dhar, N., Cole, S. T. & Chen, J. M. *Mycobacterium tuberculosis* EspK Has Active but Distinct Roles in the Secretion of EsxA and EspB. J Bacteriol 204, e0006022, (2022).

60 Orme, I. M. & Basaraba, R. J. The formation of the granuloma in tuberculosis infection. Semin Immunol 26, 601–609, (2014).

61 Wells, G. et al. A high-resolution 3D atlas of the spectrum of tuberculous and COVID-19 lung lesions. EMBO Mol Med 14, e16283, (2022).

62 Barry, C. E. Interpreting cell wall’virulence factors’ of *Mycobacterium tuberculosis*. Trends Microbiol 9, 237–241., (2001).

63 Niederweis, M., Danilchanka, O., Huff, J., Hoffmann, C. & Engelhardt, H. Mycobacterial outer membranes: in search of proteins. Trends Microbiol 18, 109–116, (2010).

64 Raman, S., et al. *Mycobacterium tuberculosis* SigM positively regulates Esx secreted protein and nonribosomal peptide synthetase genes and down regulates virulence-associated surface lipid synthesis. J Bacteriol 188, 8460–8468, (2006).

65 Walburger, A. et al. Protein kinase G from pathogenic mycobacteria promotes survival within macrophages. Science 304, 1800–1804, (2004).

66 Xander, C., Rajagopalan, S., Jacobs, W. R., Jr. & Braunstein, M. The SapM phosphatase can arrest phagosome maturation in an ESX-1 independent manner in *Mycobacterium tuberculosis* and BCG. Infect Immun 92, e0021724, (2024).

67 Abdallah, A. M. et al. PPE and PE_PGRS proteins of *Mycobacterium marinum* are transported via the type VII secretion system ESX-5. Mol Microbiol 73, 329–340, (2009).

68 Dartois, V. & Dick, T. Therapeutic developments for tuberculosis and nontuberculous mycobacterial lung disease. Nat Rev Drug Discov 23, 381–403, (2024).

69 Portal-Celhay, C., et al. *Mycobacterium tuberculosis* EsxH inhibits ESCRT-dependent CD4+ T-cell activation. Nat Microbiol 2, 16232, (2016).

70 Ates, L. S. et al. Essential Role of the ESX-5 Secretion System in Outer Membrane Permeability of Pathogenic Mycobacteria. PLoS Genet 11, e1005190, (2015).

