## Supplementary material for "Master control of protein secretion by *Mycobacterium tuberculosis*": Complete Supplement

### Supplementary Figures

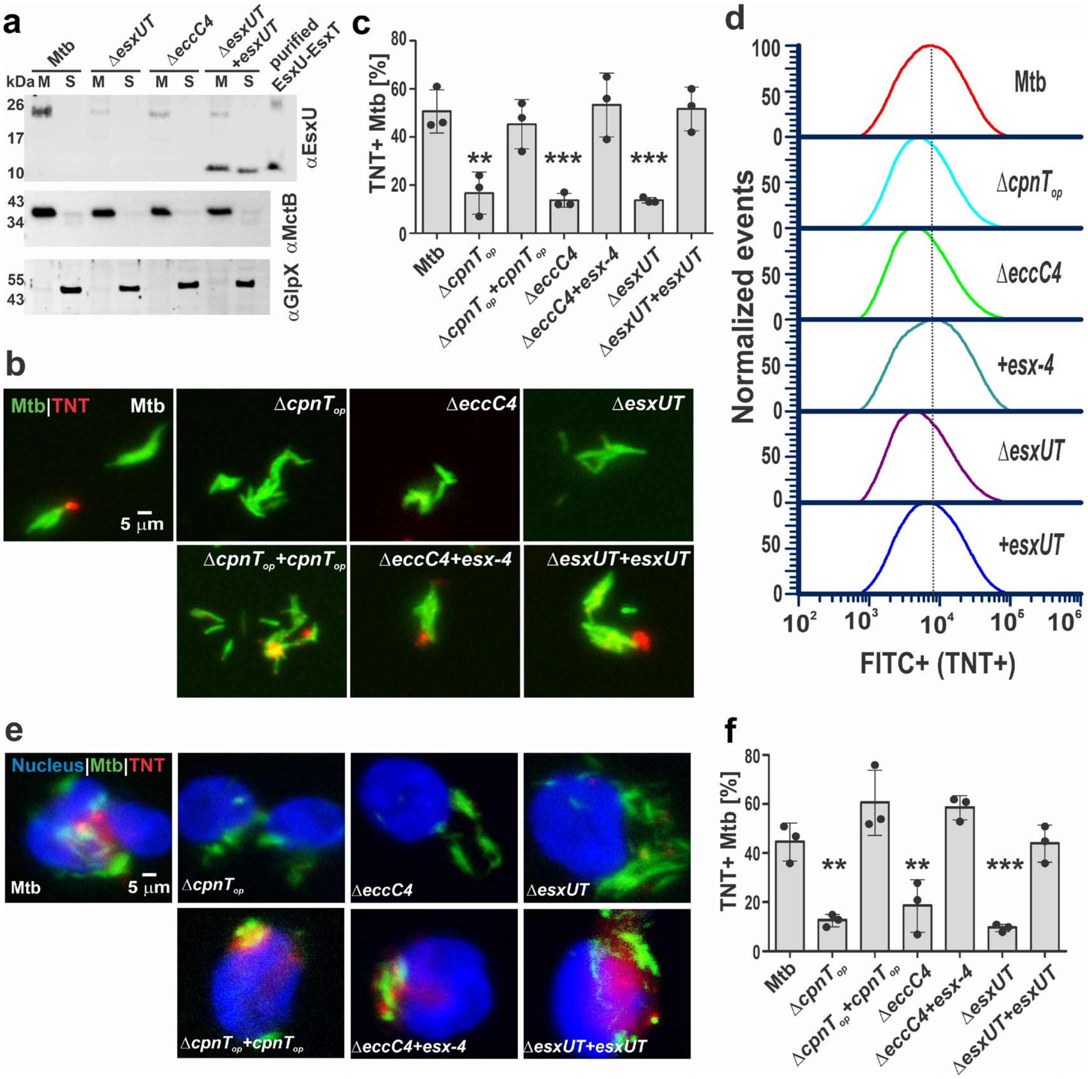

**Figure S1. EsxUT is required for TNT surface accessibility on the surface of *M. tuberculosis* and its secretion into the cytosol of infected macrophages.**

**a.** Subcellular localization of EsxU in the indicated *Mtb* strains using an anti-EsxU antiserum. The *Mtb* strain Δ*eccC4* lacks the EccC4 ATPase, which is essential for the activity of the ESX-4 system. MctB and GlpX were used as marker proteins for the membrane and cytosolic fractions, respectively. 100 ng purified EsxU-EsxT were loaded. **b.** Detection of surface accessible TNT in Mtb by fluorescence microscopy. The indicated Mtb strains were metabolically labeled with DMN-Trehalose (green) and stained with an anti-TNT antiserum (red). Scale bar = 5 µm. **c.** Quantification of surface-accessible TNT in Mtb from images shown in Fig. 1b. Strains were scored positive for TNT relative to Mtb Δ$cpnT_{op}$ (n=3). **d.** Surface accessibility of TNT in 50,000 Mtb cells by flow cytometry using anti-TNT antiserum. The peak fluorescence for wt Mtb is indicated by a dashed line. **e.** Analysis of TNT secretion into the cytosol of THP-1 macrophages infected with DMN-Trehalose labelled Mtb strains (green) at an MOI of 10:1. The cells were fixed, permeabilized with 0.2% Triton X-100 and stained using anti-TNT antiserum (red). The nuclei of the macrophages were stained with DAPI (blue). Scale bar = 5 µm. **f.** Quantification of TNT secretion into the cytosol of infected macrophages in images shown in Fig. 1e, relative to Mtb Δ$cpnT_{op}$ (n=3). All quantitative data (c, f) are represented as mean ± standard deviation (SD). Asterisks indicate significant differences (** $p \leq 0.01$, *** $p \leq 0.001$ using one-way ANOVA with Dunnett's correction) compared to wt Mtb.

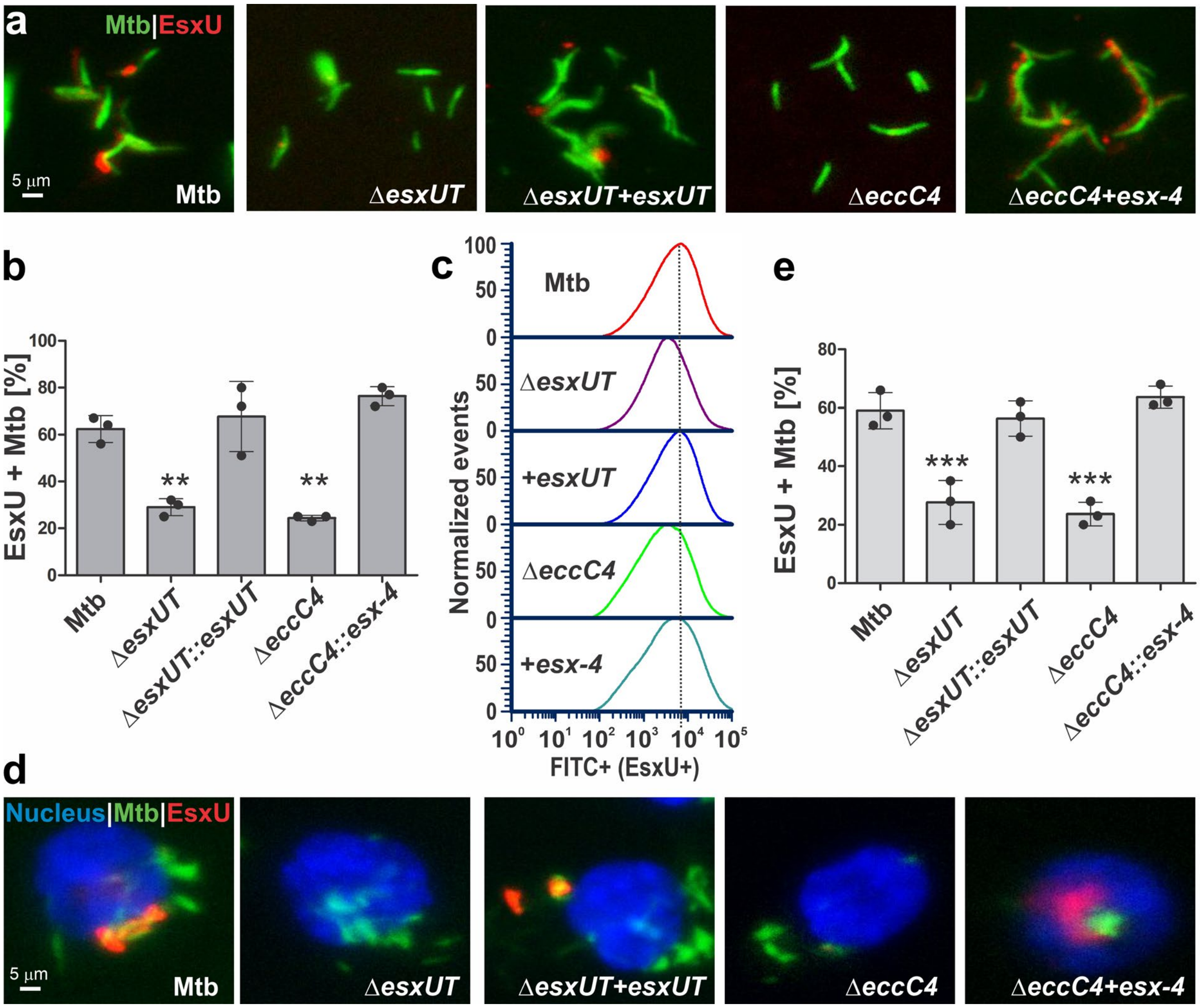

**Fig. S2: EsxU is a surface protein of *Mtb* and is secreted into the cytosol of infected macrophages in an ESX-4 dependent manner.**

**a.** Detection of surface-accessible EsxU in Mtb using fluorescence microscopy. The indicated Mtb strains were metabolically labeled with DMN-Trehalose (green) and stained with an anti-EsxU antiserum (red). The Mtb strain Δ*eccC4* lacks the EccC4 ATPase, which is essential for the activity of the ESX-4 system. Scale bar = 5 µm. **b.** Quantification of surface accessible EsxU in Mtb in the images shown in Fig. 2a (n=3). Strains were scored positive for EsxU relative to Mtb Δ*esxTU*. **c**. Surface accessibility of EsxU in 50,000 Mtb cells by flow cytometry using an anti-EsxU antiserum. The peak fluorescence for wt Mtb is indicated by a dashed line. **d**. Analysis of EsxU secretion into the cytosol of THP-1 macrophages infected with DMN-Trehalose labelled Mtb strains (green) at an MOI of 10:1. The cells were fixed, permeabilized with 0.2% Triton X-100 and stained using anti-EsxU antiserum (red). The nuclei of the macrophages were stained with DAPI (blue). Scale bar = 5 µm. **e.** Quantification of EsxU secretion into the cytosol of infected macrophages in images shown in Fig. 1d, relative to Mtb Δ*esxTU* (n=3). All quantitative data (b, e) are represented as mean ± standard deviation (SD). Asterisks indicate significant differences (** $p \leq 0.01$, *** $p \leq 0.001$ calculated using one-way ANOVA with Dunnett's correction) in comparison to wt Mtb.

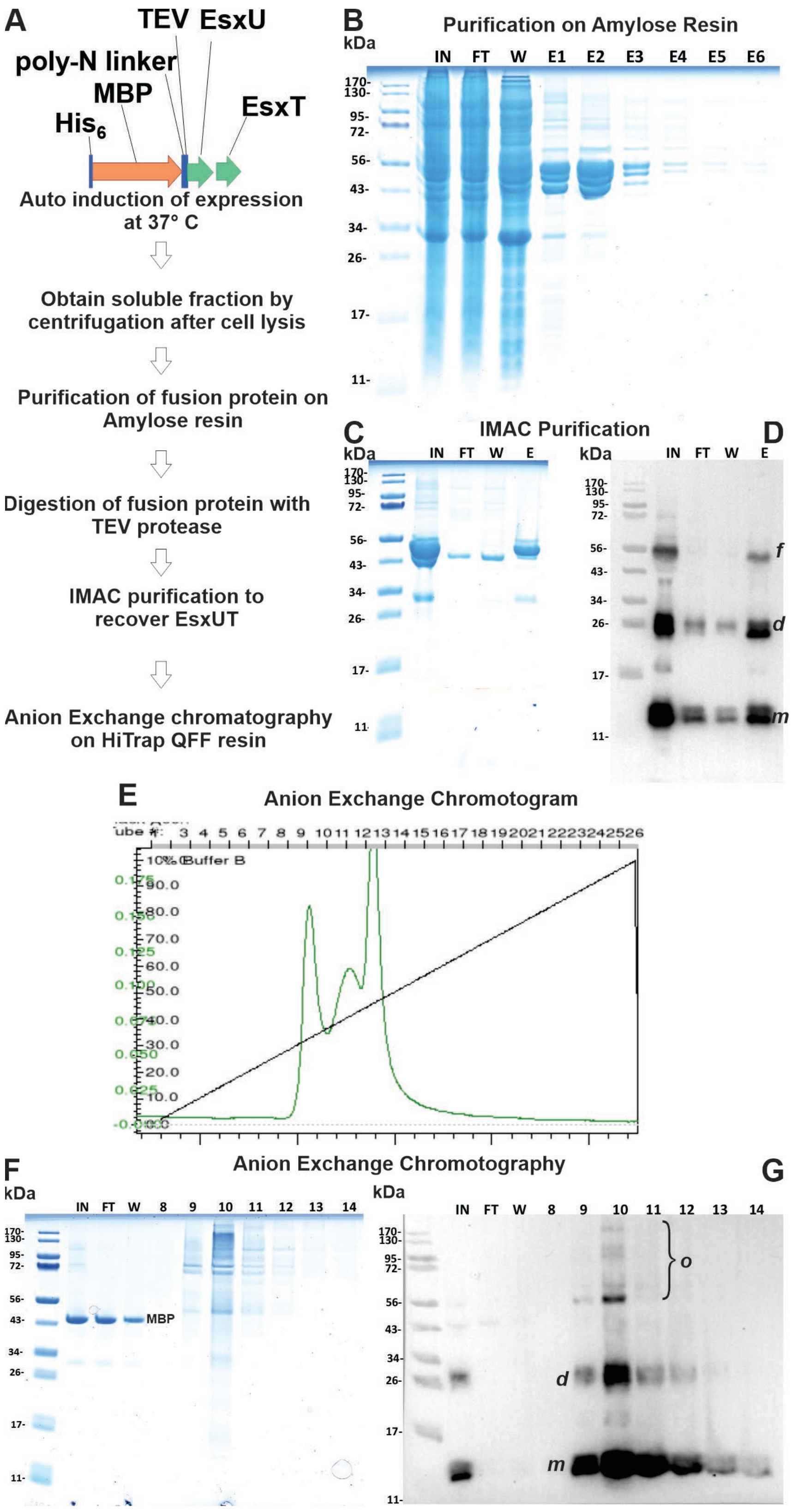

**Figure S3. Purification of the *M. tuberculosis* EsxUT complex from *E. coli*.**
**A.** Schemes of MBP-EsxUT fusion protein construct and purification steps.

**B.** SDS-PAGE of purification of MBP-EsxUT fusion protein on Amylose resin. Lanes: IN – input, soluble fraction after cell lysis and centrifugation at 50 k *g*; FT – flow-through; W – wash with column buffer without D-maltose; E1-E6 – elutions with 20 mM D-maltose buffer. 5 µg of protein was loaded per lane. Molecular weight protein standard in kDa is on the left.

**C and D.** IMAC purification of EsxUT after TEV protease treatment. Samples were analyzed on polyacrylamide gel (C) and Western blot (D) stained with EsxU antiserum. Lanes: IN – input, sample after incubation with TEV protease; FT – flow-through; W – wash with 20 mM imidazole; E – elution with 500 mM imidazole. Cleaved EsxUT (marked ***d*** for dimer, and ***m*** for monomer) was collected from FT and W fractions of the purification step. Elution fraction was discarded since it contains $His_6$-MBP-EsxUT fusion protein (marked ***f***). 5 µg of protein was loaded per lane.

**E.** Chromatogram of anion exchange purification of EsxUT on HiTrap QFF resin. EsxUT was eluted by applying 150 mM to 2 M NaCl gradient (black line). Green line is Absorbance at 280 nm. EsxUT elutes at 570 mM NaCl.

Fractions after anion exchange were loaded on polyacrylamide gel stained with Coomassie blue (**F**) and Western blot (**G**) stained with EsxU antiserum. Lanes: IN – input, sample after IMAC; FT – flow-through collected during column loading; W – column wash with 150 mM NaCl before elution with 0.15 – 2 M NaCl gradient; numbers 8-14 – elution fractions corresponding to peaks in **E**. MBP –is marked on the gel and elutes in the flow-through and washing steps at 150 mM NaCl. EsxU monomers (***m***), dimers (***d***), and high molecular weight oligomers (***o***) are visible on the Western. Same protein amount was loaded in each lane. Molecular weight protein standard in kDa is on the left.

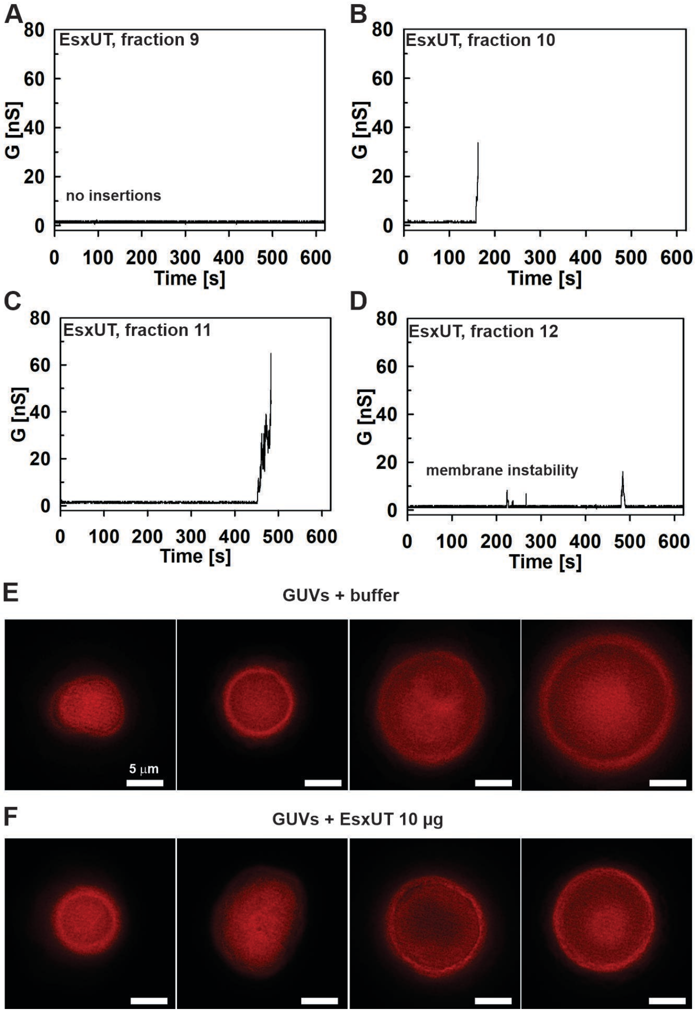

**Figure S4. Characterization of purified *M. tuberculosis* EsxUT*.***

**A - D.** Lipid bilayer experiments with EsxUT from Anion Exchange purification, Fig. SX. Current traces of individual fractions of Anion Exchange in diphytanoyl phosphatidylcholine (DPhPC) membranes are shown. The experiments were performed in 1M KCl, 25 mM sodium phosphate, pH 7.4 at 10 mV applied potential. A total of 7 membranes with fraction 9 (**A**) was recorded. No insertions were observed when 1 – 10 µg of protein was present. Five membranes with fraction 10 (**B**) was recorded with 0.88 – 15 µg of protein present. In only one membrane several fast insertions were observed and the membrane broke immediately after insertions (**B**). Similar result was obtained with fraction 11 (**C**) where one membrane out of seven resulted in fast insertions followed by membrane breakage (1 – 10 µg of protein was present). Panel **D** shows experiment with fraction 12 (1 – 10 µg of protein) where one membrane out of 7 exhibited instability.

**E.** Giant unilamellar vesicles (GUVs) treated with FM464X dye to label dimyristoyl phosphocholine (DMPC) lipid membrane. GUVs were incubated for 30 min in buffer containing 25 mM sodium phosphate, 150 mM NaCl, pH 7.0. Representative micrographs are shown. The white bar is 5 µm.

**F**. Giant unilamellar vesicles (GUVs) stained with FM464X dye. GUVs were incubated for 30 min with 10 µg of purified EsxUT in a buffer containing 25 mM sodium phosphate, 150 mM NaCl, pH 7.0. Representative micrographs are shown. The scale bar is 5 µm.

#### Co-expression of HA-EsxU/EsxT-8xHis and Myc-EsxF/EsxE-FLAG

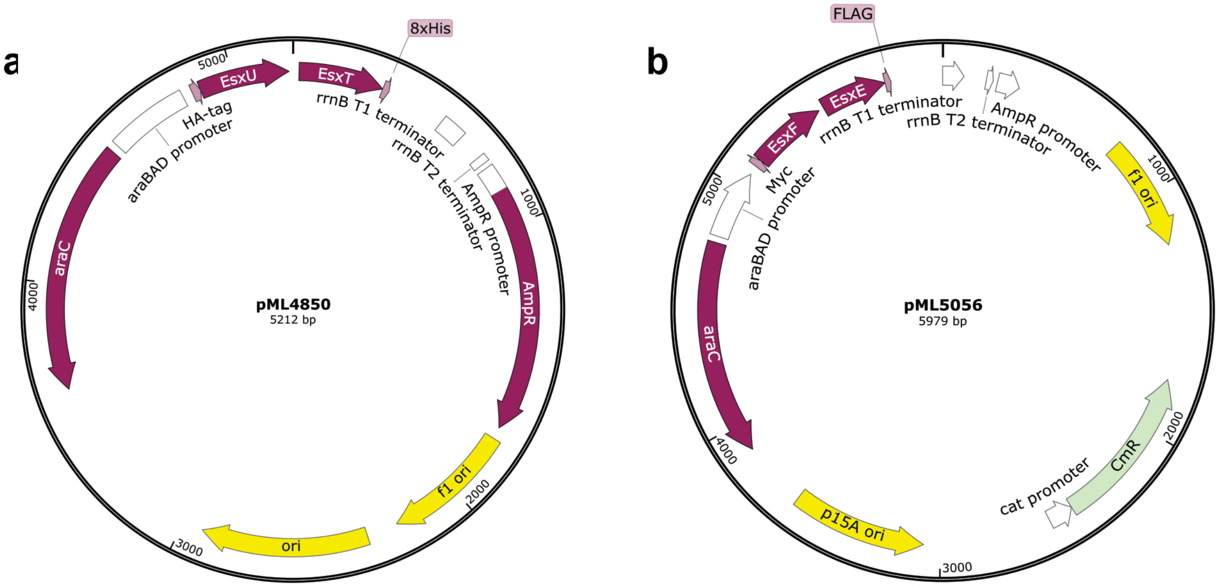

#### Co-expression of 8xHis-EsxF/EsxE-FLAG and HA-EsxU/EsxT-Myc

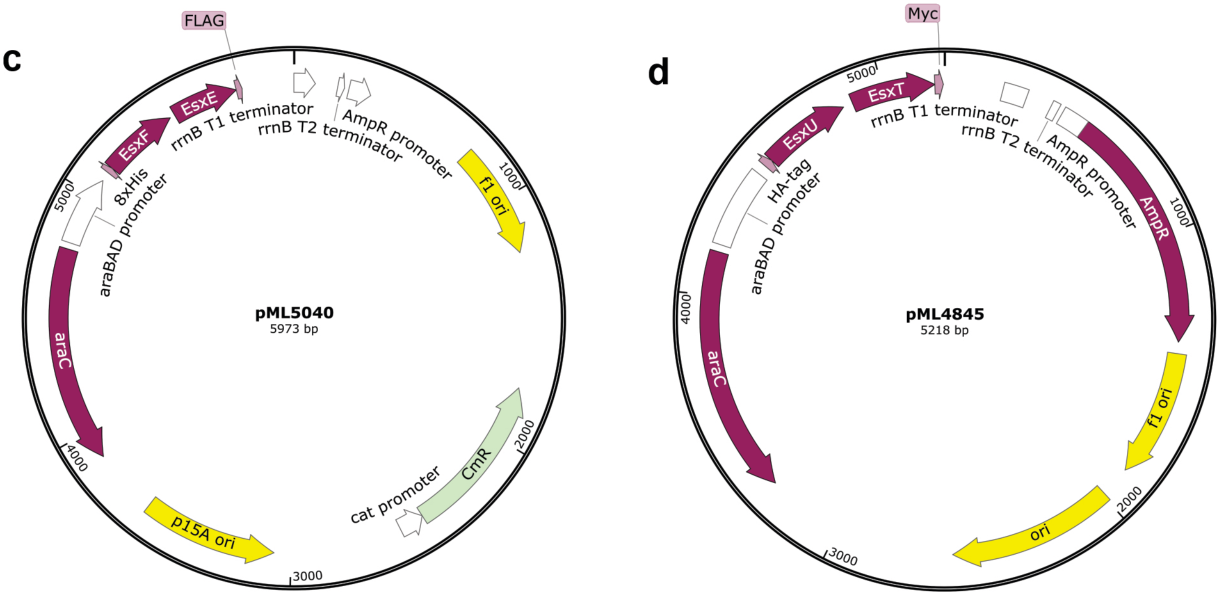

**Fig. S5. Co-expression vectors to examine the interactions of small Esx proteins in *E. coli*.**
The genomic regions of EsxU-EsxT and EsxF-EsxE were amplified from Mtb wt using specific primers containing the 1 kDa tag sequences (Table S3). The tagged sequences were inserted in the respective vectors to obtain: **a.** pML4850 expressing HA-EsxU-EsxT-8xHis; **b.** pML5056 expressing Myc-EsxF-EsxE-FLAG; **c.** pML5040 expressing 8xHis-EsxF-EsxE-FLAG; and **d.** pML4845 expressing HA-EsxU-EsxT-Myc.

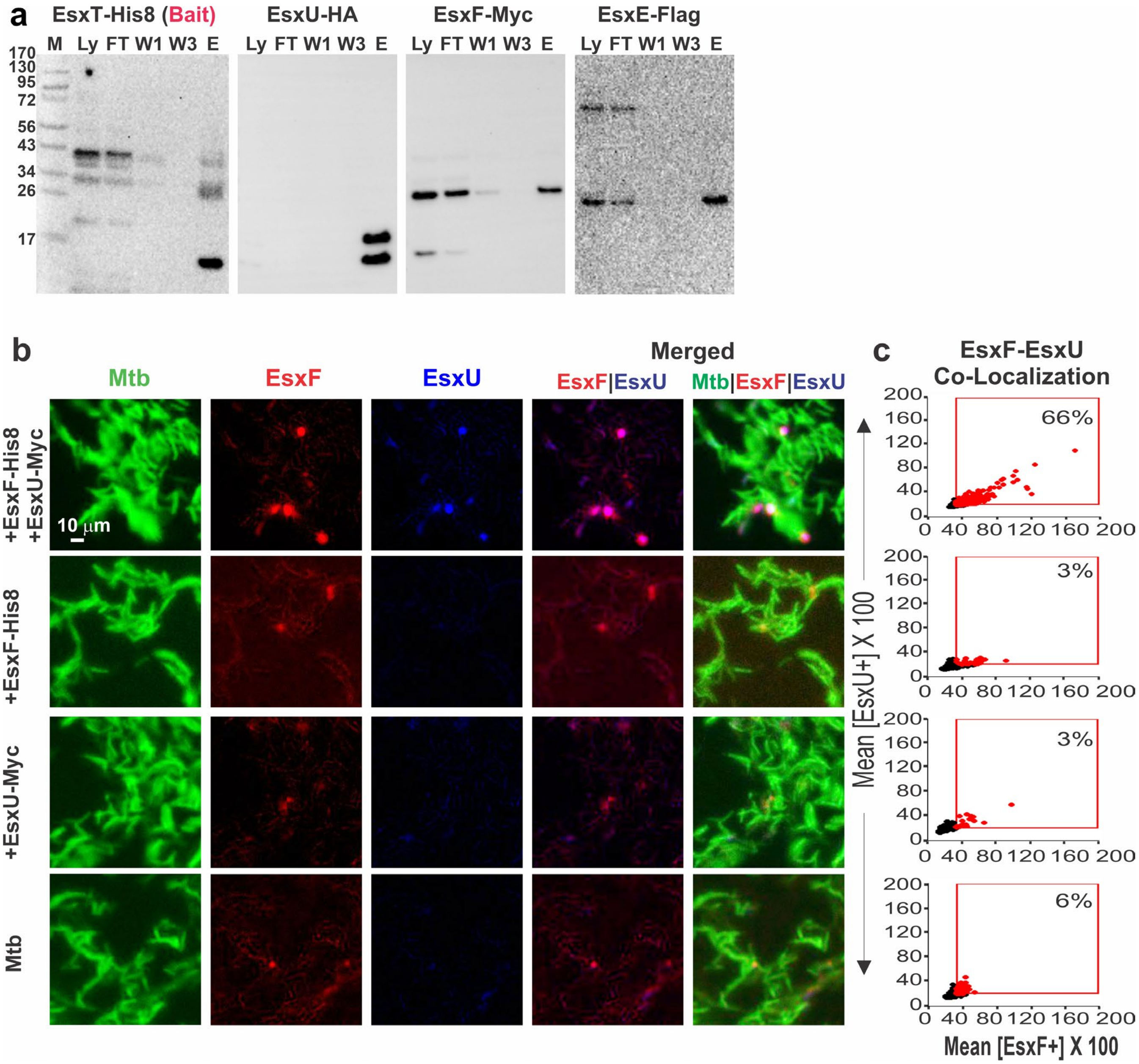

**Fig. S6. EsxEF and EsxUT interact and co-localize on the cell surface of *M. tuberculosis*.**

**a.** Pull-down assay: The bait protein EsxT$_{His8}$ in a lysate (Ly) of *E. coli* producing $_{HA}$EsxU/EsxT$_{His8}$ and $_{Myc}$EsxF/EsxE$_{FLAG}$ was captured on a Ni(II) affinity column. The 10-fold diluted cell lysate (Ly), the flow through (FT), buffers of wash one (W1) and wash three (W3) and the sample eluted with imidazol (E) were analyzed in Western blots using tag-specific antibodies. **b.** The surface accessibility of tagged EsxF and EsxU proteins in Mtb strains producing either $_{His8}$EsxF-EsxE or $_{Myc}$EsxU-EsxT or both Esx pairs together. Strains were scored positive for tagged EsxF or EsxU relative to untagged wt Mtb. The indicated Mtb strains were metabolically labeled with DMN-trehalose (green) and stained with anti-His (red) and anti-Myc (blue) Alexa fluor-conjugated antibodies. Co-localization of EsxF and EsxU in these Mtb strains was visualized by fluorescence micoscopy showing the merged images and quantified as a percentage of positive cells (**c**).

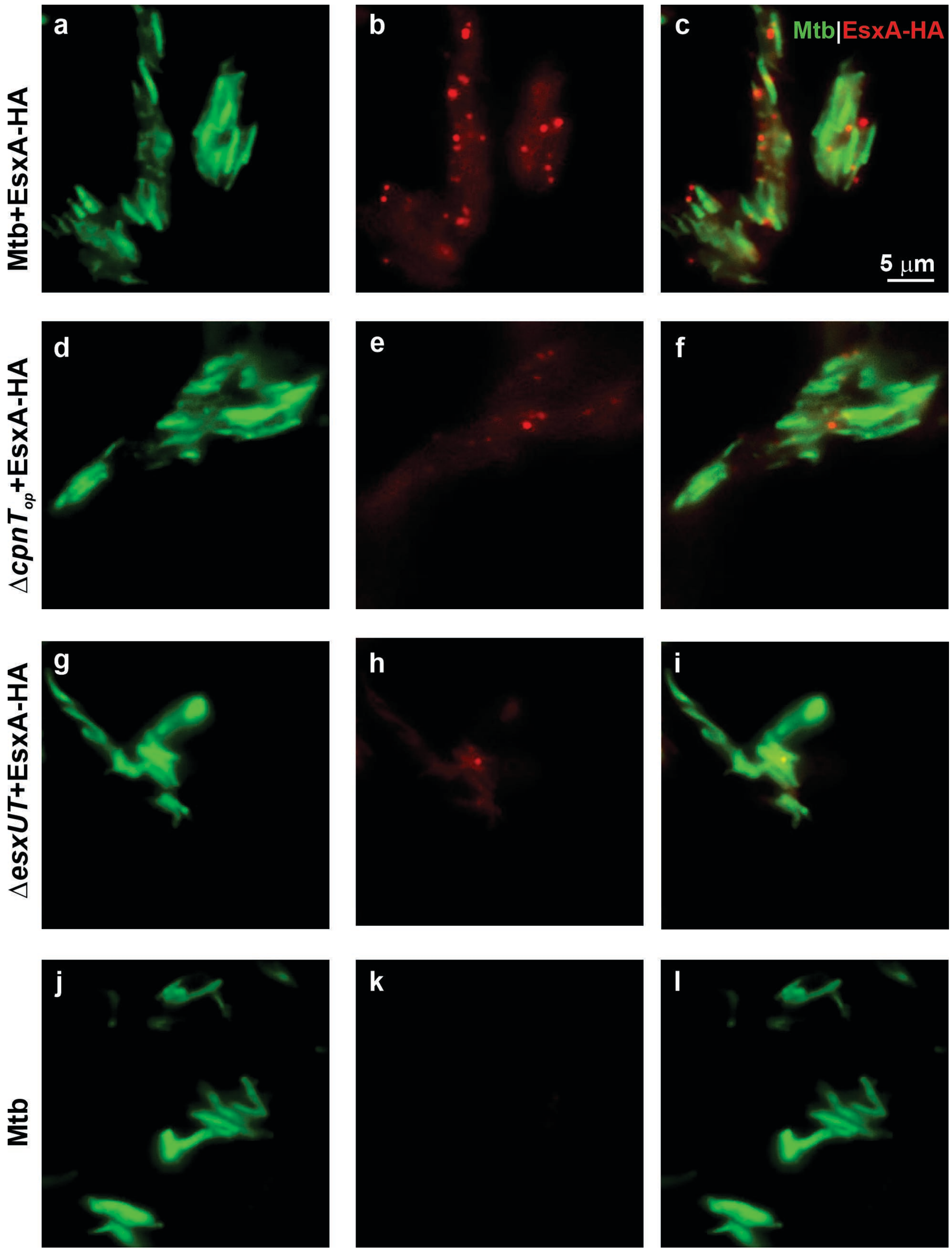

**Fig. S7: Role of EsxEF-EsxUT in surface exposure of EsxA in *M. tuberculosis*.**
These high resolution images of the experiment shown in Fig. 5A were acquired using a fluorescence microscope with a 100X oil objective phase. The parent strain *M. tuberculosis* mc$^2$6206 (Mtb) and derivative strains were metabolically labeled with DMN-trehalose (green) and were probed for HA-tagged EsxA using a monoclonal HA-tag antibody and an Alexa fluor 594-conjugated secondary antibody (red). HA tagged EsxA levels in: **a-c.** Mtb mc$^2$6206, **d-f.** Mtb Δ*cpnT*$_{op}$, and **g-i.** Mtb Δ*esxUT*. The fluorescence of these strains was normalized compared to Mtb mc$^2$6206, which does not produce an HA-tagged protein (**k**).

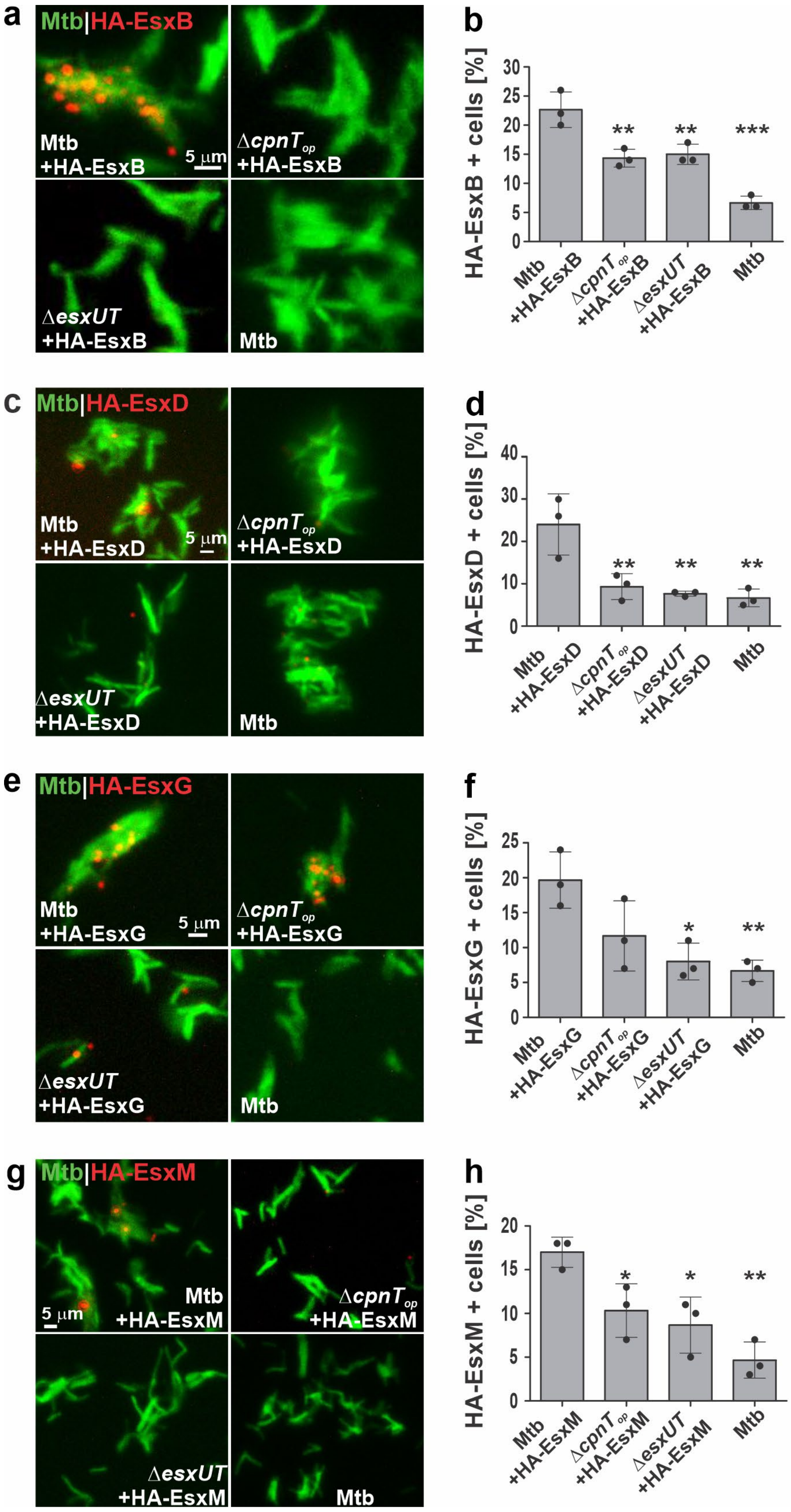
a
Mtb|HA-EsxB
Mtb
+HA-EsxB
5 µm
ΔcpnT_op
+HA-EsxB
ΔesxUT
+HA-EsxB
Mtb
b
HA-EsxB + cells [%]
0
5
10
15
20
25
30
**
**
***
Mtb +HA-EsxB
ΔcpnT_op +HA-EsxB
ΔesxUT +HA-EsxB
Mtb
c
Mtb|HA-EsxD
Mtb
+HA-EsxD
5 µm
ΔcpnT_op
+HA-EsxD
ΔesxUT
+HA-EsxD
Mtb
d
HA-EsxD + cells [%]
0
10
20
30
40
**
**
**
Mtb +HA-EsxD
ΔcpnT_op +HA-EsxD
ΔesxUT +HA-EsxD
Mtb
e
Mtb|HA-EsxG
Mtb
+HA-EsxG
5 µm
ΔcpnT_op
+HA-EsxG
ΔesxUT
+HA-EsxG
Mtb
f
HA-EsxG + cells [%]
0
5
10
15
20
25
*
**
Mtb +HA-EsxG
ΔcpnT_op +HA-EsxG
ΔesxUT +HA-EsxG
Mtb
g
Mtb|HA-EsxM
5 µm
Mtb
+HA-EsxM
ΔcpnT_op
+HA-EsxM
ΔesxUT
+HA-EsxM
Mtb
h
HA-EsxM + cells [%]
0
5
10
15
20
*
*
**
Mtb +HA-EsxM
ΔcpnT_op +HA-EsxM
ΔesxUT +HA-EsxM
Mtb

**Fig. S8: Role of EsxEF-EsxUT in surface exposure of EsxB and ESX-associated EsxB homologs in *M. tuberculosis*.**

Fluorescence microscope images of surface accessible HA-tagged: **a.** EsxB, **c.** EsxD, **e.** EsxG, and **g.** EsxM in the parent strain *M. tuberculosis* $mc^2$6206 (Mtb) and derivative strains. DMN-Trehalose stained Mtb strains (green) were probed for the HA-tagged small Esx proteins (red). Scale bar = 5 µm. Quantification of surface exposed HA tagged: **b.** EsxB, **d.** EsxD, **f.** EsxG, and **h.** EsxM in comparison to the untagged WT cells, determined from Cytation 5 analysis (n=3). Data are represented as mean ± SD and asterisks indicate significant differences (*p value ≤ 0.05, **p value ≤ 0.01, ***p value ≤ 0.001 calculated using the one-way ANOVA with Dunnett's correction) in comparison to the tagged WT strain.

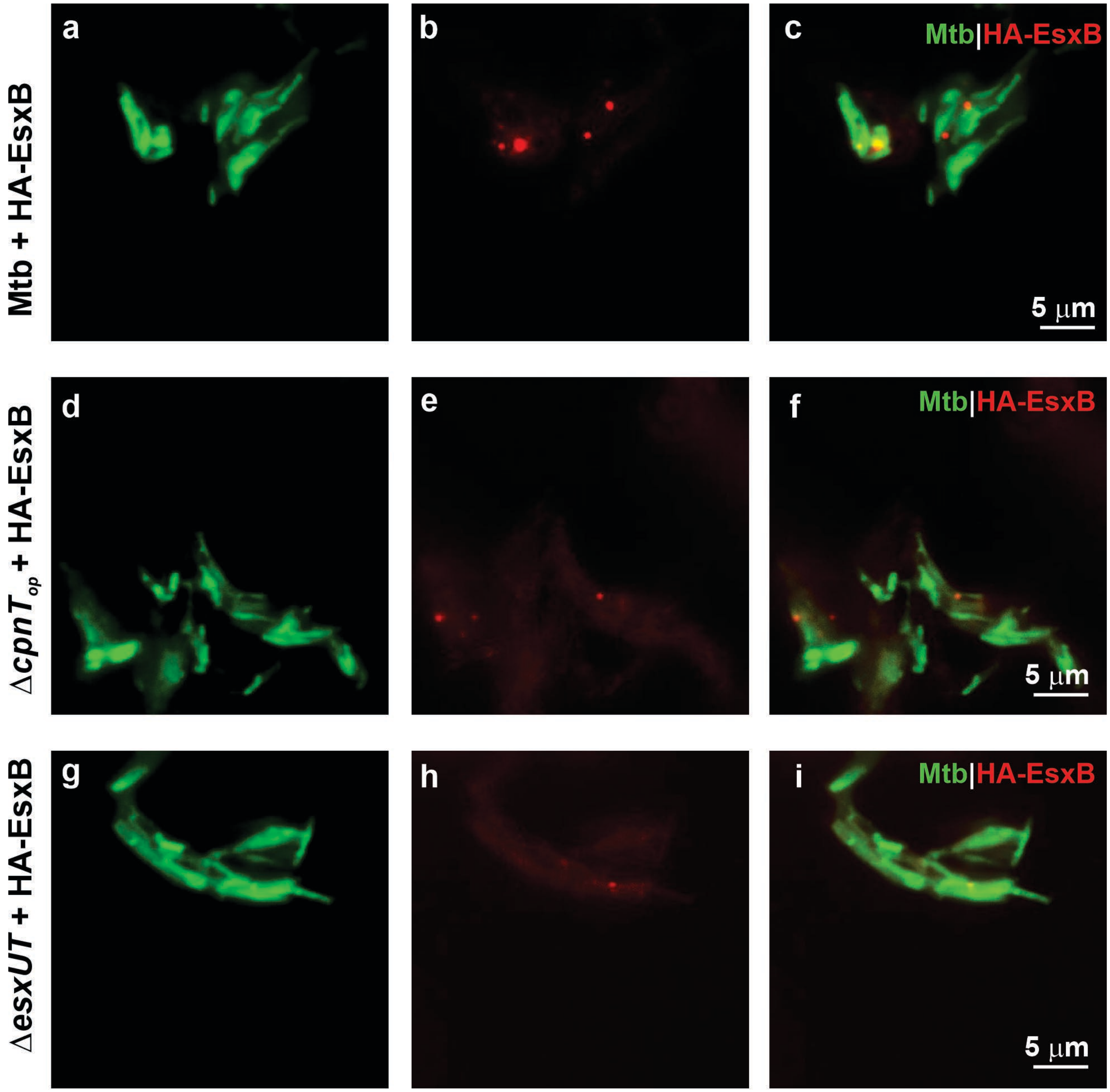

**Fig. S9: Role of EsxEF-EsxUT in surface exposure of EsxB in *M. tuberculosis*.**
These high resolution images of the experiment shown in Fig. S8A were acquired using a fluorescence microscope with a 100X oil objective phase. DMN-Trehalose stained Mtb strains (green) were probed for HA-tagged EsxB (red) in the parent strain *M. tuberculosis* mc$^2$6206 (Mtb) and derivative strains. HA tagged EsxB levels in: **a-c.** Mtb mc$^2$6206, **d-f.** Mtb Δ*cpnTop*, and **g-i.** Mtb Δ*esxUT*. The HA fluorescence levels in the strains were normalized compared to the untagged Mtb mc$^2$6206 strain (Fig. S8k).

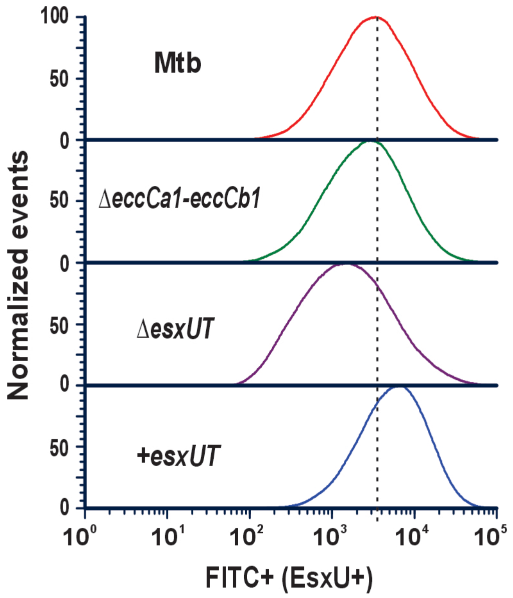

**Fig. S10: Surface exposure of EsxU in Mtb does not depend on the ESX-1 system.**
Surface-accessible EsxU in the parent strain *M. tuberculosis* mc$^2$6206 (Mtb) and derivative strains were identified using anti-EsxU antiserum and Alexa fluor 488 conjugated secondary antibody by flow cytometry. The peak value of fluorescence for Mtb mc$^2$6206 is marked by a dashed line in the histogram.

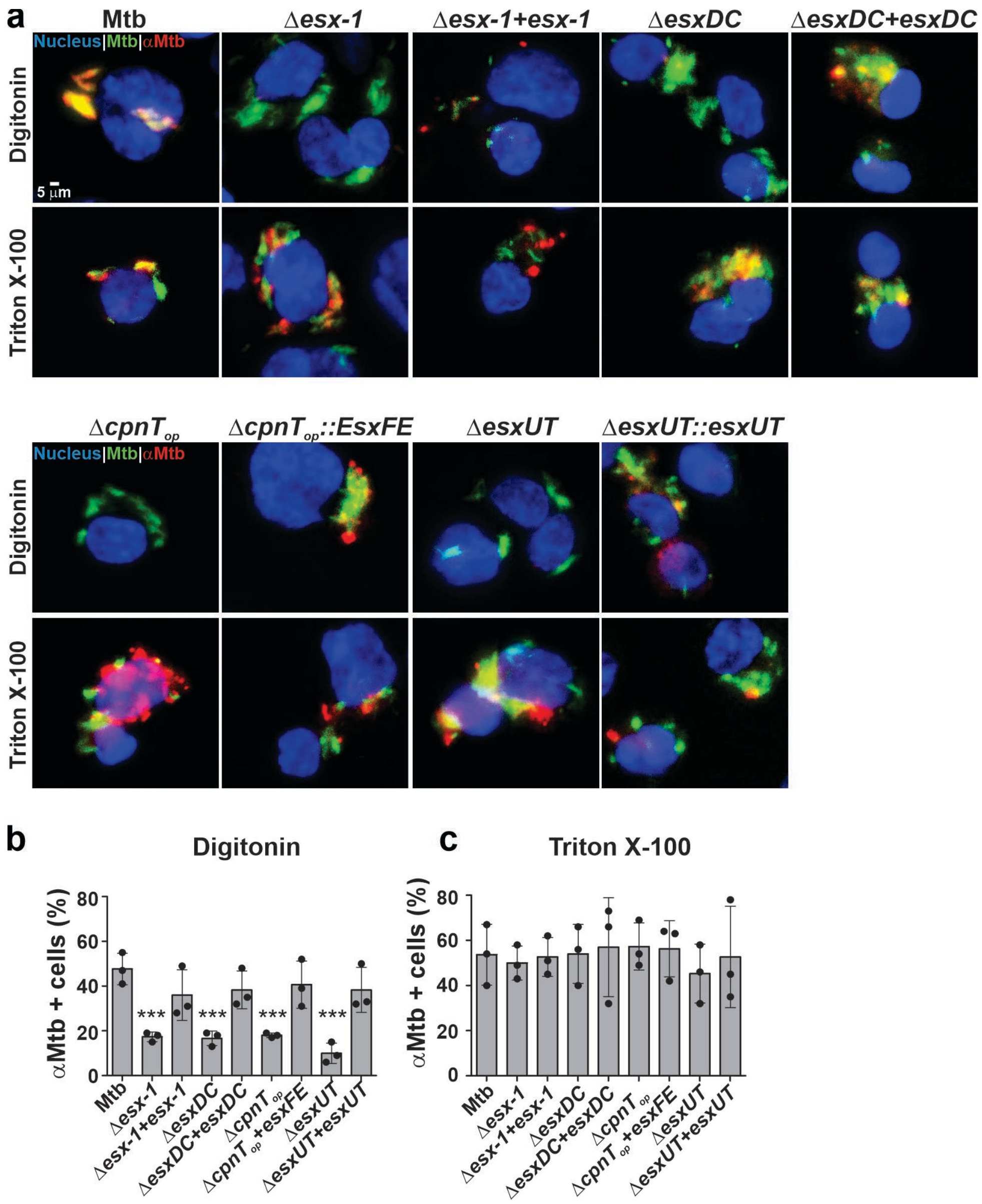

**Fig. S11: EsxEF and the Esx effector proteins of the ESX-1, ESX-2 and ESX-4 systems are required for phagosomal permeabilization by *M. tuberculosis*.**

**a.** Determination of phagosomal permeabilization in macrophages using Mtb specific antibodies. The DMN-trehalose labeled parent strain *M. tuberculosis* mc$^2$6206 (Mtb) and derivative strains were used to infect the THP-1 macrophages at an MOI of 10:1. The macrophages were fixed after 48 h of infection, permeabilized either with digitonin which facilitates the access of antibodies in an antiserum against the tuberculin purified protein derivative (PPD) (anti-Mtb antibody) to the cytosol of macrophages or with Triton X-100 to enable the access of antibodies to the phagosomal compartment {Pajuelo, 2021 #64035}. Cells were incubated with an anti-Mtb antibody (αMtb) and an Alexa Fluor-594 secondary antibody (red). The nuclei of the infected macrophages were stained with DAPI and analyzed using Cytation 5. Scale bar = 5 µm. **b, c.** Quantification of infected macrophages positive for Mtb proteins in comparison to the *eccCa1-eccCb1* deletion mutant when permeabilized with: **b.** Digitonin, and **c.** Triton X-100, (n=3). Data are represented as mean ± SD and asterisks indicate significant differences (***p value ≤ 0.001 calculated using the one-way ANOVA with Dunnett's correction) in comparison to Mtb mc$^2$6206 strain.

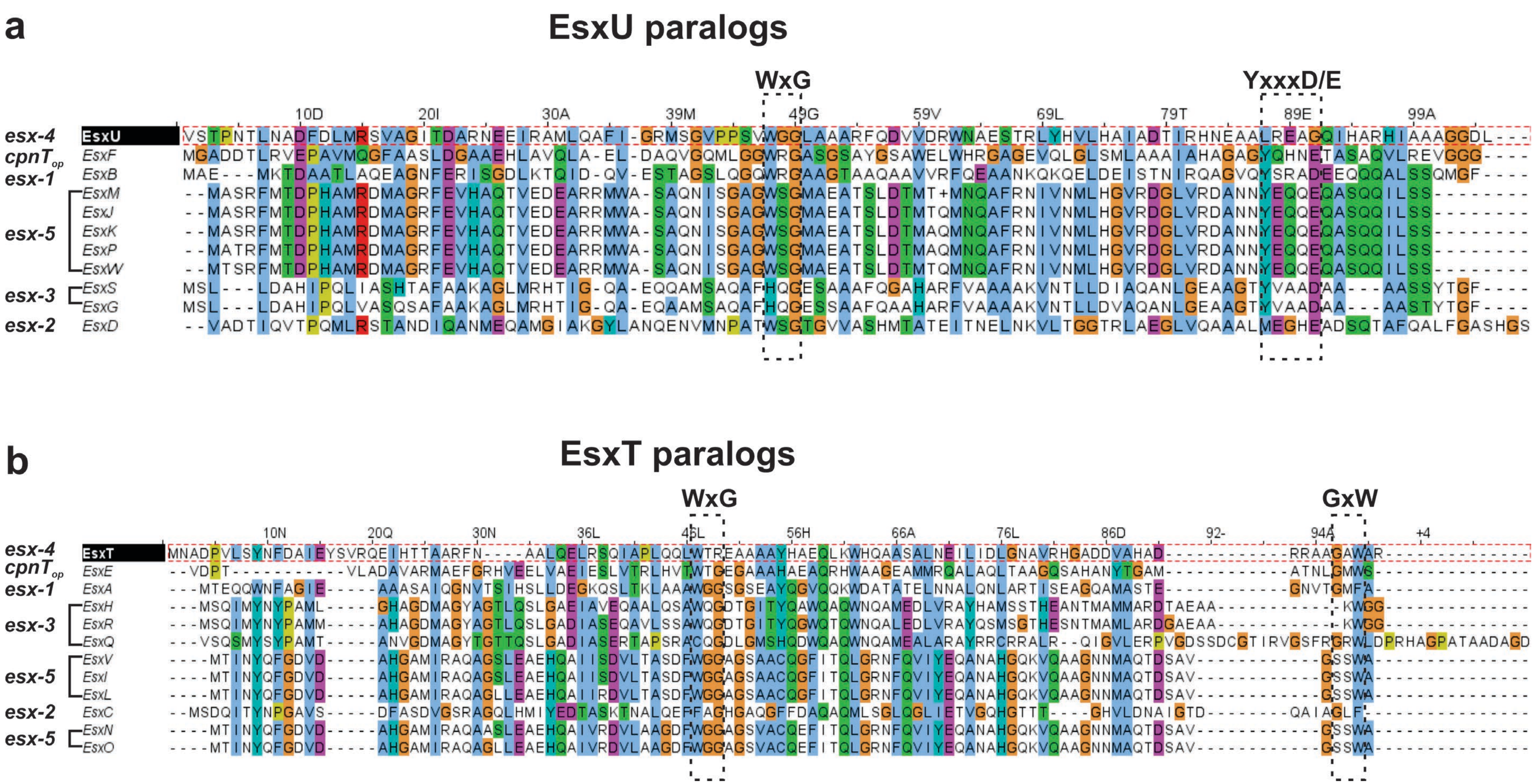

**Fig. S12: Sequence alignment of EsxU and EsxT paralogs**

Multiple sequence alignments of EsxU (**a**) and EsxT (**b**) paralogs were generated using Clustal Omega. The sequences were obtained from Mycobrowser (Release 5), aligned in Clustal Omega and visualized using the Jalview software. The associations of the Esx proteins with their cognate type VII secretion systems is indicated on the left. The locations of the type VII secretion signal YxxxD/E and of the WxG and GxW motifs are shown in sequence alignments.

The color codes are:

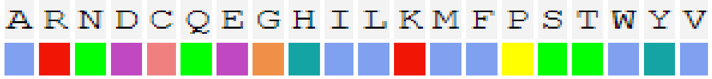

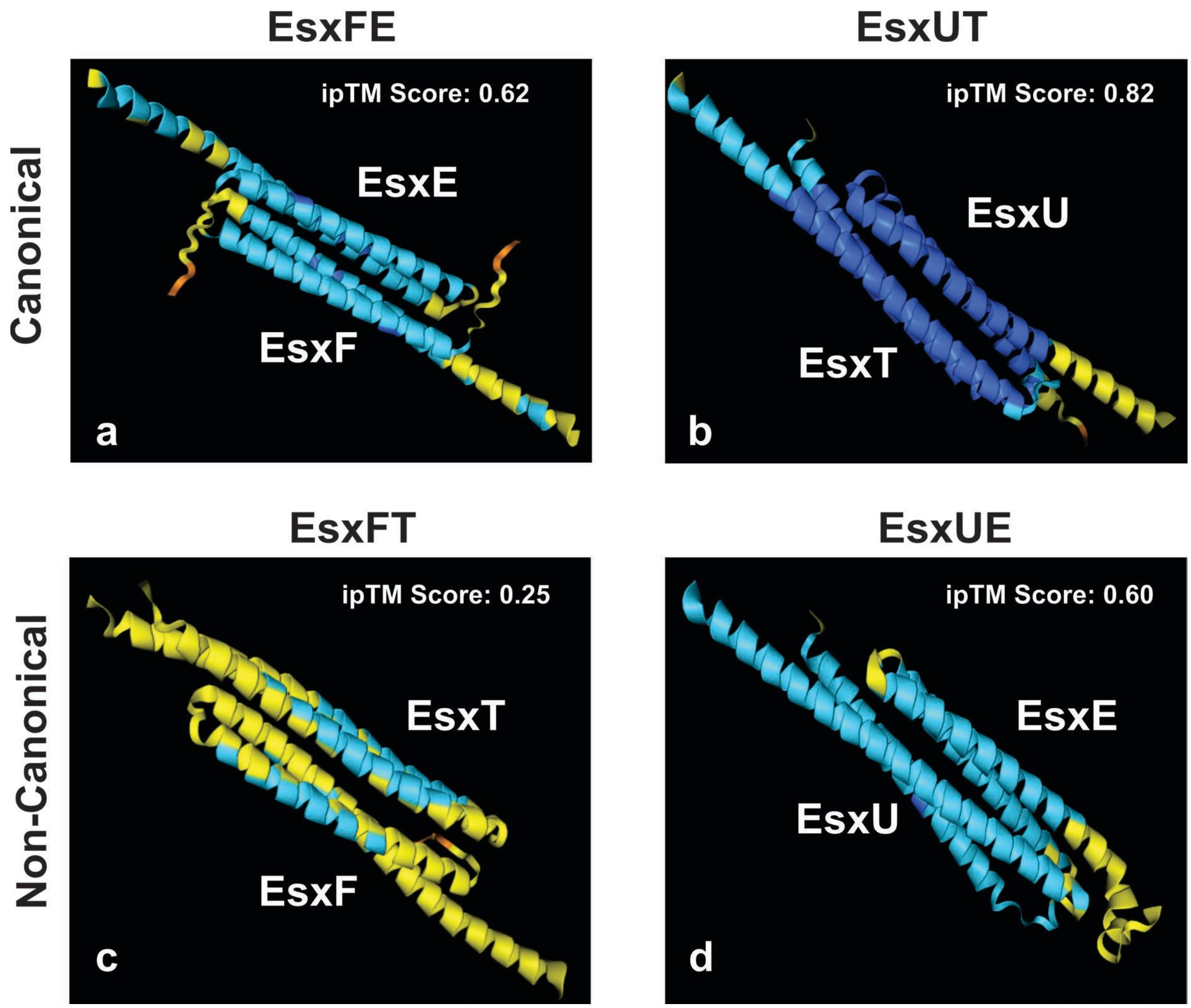

**Fig. S13: Alphafold models for the interactions of EsxEF and EsxUT of *M. tuberculosis*.** Alphafold models show the probable interaction between (**a, b**) canonical heterodimers: **a.** EsxFE and **b.** EsxUT, and (**c, d**) non-canonical heterodimers: **c.** EsxFT and **d.** EsxUE. The interface predicted template modelling (ipTM) score of each protein complex is indicated.

### Supplementary Tables

| Strain name | Relevant genotypes and description | Source or Reference |
|---|---|---|
| *M. tuberculosis* mc$^2$6206 | H37Rv derivative; Δ*leuCD* Δ*panCD*; avirulent *M. tuberculosis* (BSL-2) | (Sampson et al., 2004) |
| *M. tuberculosis* ML2016 | mc$^2$6206 derivative; Δ*esxF-esxE-cpnT-ift::loxP* (referred to as Δ*cpnT* operon) | (Tak et al., 2021) |
| *M. tuberculosis* ML2648 | ML2016 derivative; L5 attB::pML3964 | (Pajuelo et al., 2021) |
| *M. tuberculosis* ML2690 | ML2769 derivative; Δ*eccC4::loxP* | (Pajuelo et al., 2021) |
| *M. tuberculosis* ML2696 | ML2690/cosmid I60 (contains *esx-4* operon) | (Pajuelo et al., 2021) |
| *M. tuberculosis* ML3006 | mc$^2$6206 derivative; Δ*esxUT::hyg*$^R$ | This study |
| *M. tuberculosis* ML3002 | ML3006 derivative; Δ*esxUT::loxP* | This study |
| *M. tuberculosis* ML3003 | ML3002 derivative; L5 attB::pML4328 | This study |
| *M. tuberculosis* ML2636 | ML2016 derivative; pML3938; L5 attB::pML3954 | (Tak et al., 2021) |
| *M. tuberculosis* ML2640 | ML2016 derivative; pML3938; L5 attB::pML3958 | (Tak et al., 2021) |
| *M. tuberculosis* ML3007 | mc$^2$6206 derivative; Ms6 attB::pML5084 | This study |
| *M. tuberculosis* ML3008 | mc$^2$6206 derivative; L5 attB::pML4862 | This study |
| *M. tuberculosis* ML3009 | mc$^2$6206 derivative; Ms6 attB::pML5084; L5 attB::pML4862 | This study |
| *M. tuberculosis* ML3010 | mc$^2$6206 derivative; L5 attB::pML4863 | This study |
| *M. tuberculosis* ML3011 | ML2016 derivative; L5 attB::pML4863 | This study |
| *M. tuberculosis* ML3012 | ML3002 derivative; L5 attB::pML4863 | This study |
| *M. tuberculosis* ML3013 | mc$^2$6206 derivative; L5 attB::pML4864 | This study |
| *M. tuberculosis* ML3014 | ML2016 derivative; L5 attB::pML4864 | This study |
| *M. tuberculosis* ML3015 | ML3002 derivative; L5 attB::pML4864 | This study |
| *M. tuberculosis* ML3016 | mc$^2$6206 derivative; L5 attB::pML4854 | This study |
| *M. tuberculosis* ML3017 | ML2016 derivative; L5 attB::pML4854 | This study |
| *M. tuberculosis* ML3018 | ML3002 derivative; L5 attB::pML4854 | This study |
| *M. tuberculosis* ML3019 | mc$^2$6206 derivative; L5 attB::pML4855 | This study |
| *M. tuberculosis* ML3020 | ML2016 derivative; L5 attB::pML4855 | This study |
| *M. tuberculosis* ML3021 | ML3002 derivative; L5 attB::pML4855 | This study |

**Table S1. Strains used in this study**

| Strain name | Relevant genotypes and description | Source or Reference |
|---|---|---|
| *M. tuberculosis* ML3022 | mc$^2$6206 derivative; L5 attB::pML4860 | This study |
| *M. tuberculosis* ML3023 | ML2016 derivative; L5 attB::pML4860 | This study |
| *M. tuberculosis* ML3024 | ML3002 derivative; L5 attB::pML4860 | This study |
| *M. tuberculosis* ML3025 | mc$^2$6206 derivative; L5 attB::pML4861 | This study |
| *M. tuberculosis* ML3026 | ML2016 derivative; L5 attB::pML4861 | This study |
| *M. tuberculosis* ML3027 | ML3002 derivative; L5 attB::pML4861 | This study |
| *M. tuberculosis* ML3028 | mc$^2$6206 derivative; L5 attB::pML4869 | This study |
| *M. tuberculosis* ML3029 | ML2016 derivative; L5 attB::pML4869 | This study |
| *M. tuberculosis* ML3030 | ML3002 derivative; L5 attB::pML4869 | This study |
| *M. tuberculosis* ML3031 | mc$^2$6206 derivative; L5 attB::pML4870 | This study |
| *M. tuberculosis* ML3032 | ML2016 derivative; L5 attB::pML4870 | This study |
| *M. tuberculosis* ML3033 | ML3002 derivative; L5 attB::pML4870 | This study |
| *M. tuberculosis* ML3034 | mc$^2$6206 derivative; Δ*eccCa1-eccCb1::hyg*$^R$ | This study |
| *M. tuberculosis* ML3001 | ML3034 derivative; Δ*eccCa1-eccCb1::loxP* | This study |
| *M. tuberculosis* ML3035 | ML3001 derivative; cosmid 2F9 | This study |
| *M. tuberculosis* ML3036 | mc$^2$6206 derivative; Δ*esxDC::hyg*$^R$ | This study |
| *M. tuberculosis* ML3000 | ML3036 derivative; Δ*esxDC::loxP* | This study |
| *M. tuberculosis* ML3037 | ML3000 derivative; L5 attB::pML4852 | This study |
| *E. coli* DH5α | F- Φ80Δ*lacZ*ΔM15 *Δ(lacZYA-argF) U169 recA1 endA1 hsdR17*(rk-mk+) *phoA supE44 thi-1 gyrA96 relA1 λ-* | Invitrogen |
| *E. coli* Mach1 | F- φ80*(lac*Z)ΔM15 Δ*lac*X74 *hsd*R(rK-mK+) Δ*rec*A1398 *end*A1 *ton*A | Invitrogen |

**Table S1. Strains used in this study (continued)**

| Plasmid | Properties | Reference |
|---|---|---|
| pML2424 | pUC origin, pAL5000ts, *loxP*-$P_{smyc}$::*gfp*$_{m}^{2+}$-*hyg*-*loxP*, $P_{wmyc}$::*sacR*-*sacB;* hyg$^R$ , vector for gene deletion in mycobacteria via homologous recombination | (Mitra et al., 2017) |
| pML4806 | pML2424-derivative; hom$_{up}$-*esxU*-*esxT*- hom$_{down}$; hyg$^R$, *esxUT* deletion vector | This study |
| pML2714 | pUC origin; pAL5000ts; $P_{hsp60}$::*cre*; $P_{imyc}$::*pamcherry1m*; *aph* | (Ofer et al., 2012) |
| pML4328 | ColE1 origin; *attP*$_{L5}$; $P_{smyc}$::*esxU*-*esxT; aph*; integrative *esxU*-*esxT* expression vector | This study |
| pBAD24-sfGFPx1 | pBR322 origin; $P_{BAD}$::*sfgfpx1*; *bla*; *E.coli* replicative expression vector for Superfolder GFP | (Malagon, 2013) |
| pML4850 | pBAD24-sfGFPx1-derivative; $P_{BAD}$::HA-*esxU*-*esxT*-8xHis; *bla*; *E.coli* replicative tagged *esxU*-*esxT* expression vector | This study |
| pBAD33 | pACYC184 origin; $P_{BAD}$; *cat*; *E.coli* replicative expression vector | (Chang and Cohen, 1978) |
| pML5056 | pBAD33-derivative; $P_{BAD}$::Myc-*esxF*-*esxE*-FLAG; *cat*; *E.coli* replicative tagged *esxF*-*esxE* expression vector | This study |
| pML5040 | pBAD33-derivative; $P_{BAD}$::8xHis-*esxF*-*esxE*-FLAG; *cat*; *E.coli* replicative tagged *esxF*-*esxE* expression vector | This study |
| pML4845 | pBAD24-sfGFPx1-derivative; $P_{BAD}$::HA-*esxU*-*esxT*-Myc; *bla*; *E.coli* replicative tagged *esxU*-*esxT* expression vector | This study |
| pML4862 | ColE1 origin; *attP*$_{L5}$; $P_{smyc}$::Myc-*esxU*-*esxT*; *aph*; integrative tagged *esxU*-*esxT* expression vector | This study |
| pML5084 | ColE1 origin; *attP*$_{Ms6}$; $P_{smyc}$::8xHis-*esxF*-*esxE*; hyg$^R$; integrative tagged *esxF*-*esxE* expression vector | This study |
| pML4863 | ColE1 origin; *attP*$_{L5}$; $P_{smyc}$::HA-*esxB*-*esxA*; *aph*; integrative tagged *esxB*-*esxA* expression vector | This study |
| pML4864 | ColE1 origin; *attP*$_{L5}$; $P_{smyc}$::*esxB*-*esxA*-HA; *aph*; integrative tagged *esxB*-*esxA* expression vector | This study |
| pML4854 | ColE1 origin; *attP*$_{L5;}$ $P_{smyc}$::HA-*esxD*-*esxC*; *aph*; integrative tagged *esxD*-*esxC* expression vector | This study |
| pML4855 | ColE1 origin; *attP*$_{L5;}$ $P_{smyc}$::*esxD*-*esxC*-HA; *aph*; integrative tagged *esxD*-*esxC* expression vector | This study |
| pML4860 | ColE1 origin; *attP*$_{L5;}$ $P_{smyc}$::HA-*esxG*-*esxH*; *aph*; integrative tagged *esxG*-*esxH* expression vector | This study |
| pML4861 | ColE1 origin; *attP*$_{L5}$; $P_{smyc}$::*esxG*-*esxH*-HA; *aph*; integrative tagged *esxG*-*esxH* expression vector | This study |
| pML4869 | ColE1 origin; *attP*$_{L5}$; $P_{smyc}$::HA-*esxM*-*esxN*; *aph*; integrative tagged *esxM*-*esxN* expression vector | This study |
| pML4870 | ColE1 origin; *attP*$_{L5}$; $P_{smyc}$::*esxM*-*esxN*-HA; *aph*; integrative tagged *esxM*-*esxN* expression vector | This study |
| pML4805 | pML2424-derivative; hom$_{up}$-*esxD*-*esxC*- hom$_{down}$; hyg$^R$; *esxDC* deletion vector | This study |
| pML4852 | ColE1 origin; *attP*$_{L5;}$ $P_{smyc}$::*esxD*-*esxC*; *aph*; integrative *esxDC* expression vector | This study |
| pML4803 | pML2424-derivative; hom$_{up}$-*eccCa1*-*eccCb1*- hom$_{down}$; hyg$^R$; *eccC1* deletion vector | This study |
| Cosmid 2F9 | pYUB412 cosmid; *bla; hyg; int*$_{L5;}$ *attP*$_{L5}$;<br>coordinates of the integrated H37Rv DNA: ~4337-4369 kb contains the ***esx-1* locus** of *M. tuberculosis* H37Rv | (Pym et al., 2002) |

**Table S2. Plasmids and cosmids used in this study**

The origin of replication is denoted as “origin”. pAL5000ts denotes the temperature-sensitive origin of replication (Guilhot et al., 1992) of the pAL5000 plasmid (Labidi et al., 1992). The *bla*, $hyg^R$, *cat* and *aph* genes confer resistance to ampicillin, hygromycin, chloramphenicol and kanamycin, respectively. The site specific integration of plasmids into the chromosomal *attB* site is mediated by the $attP_{L5}$ site, facilitated by the mycobacteriophage L5 integrase gene, $int_{L5}$. The DNA fragments flanked by *loxP* sites are excised by Cre, the site-specific recombinase.

| Oligonucleotide | Sequence (5' to 3') | Purpose |
|---|---|---|
| SpeI-Up-esxU | TGACTAGTCAGATCGCGGAGCGACTGGTGTTG | pML4806 |
| EsxUT-2 | ATTTAAATCCTCGTCTGGTTCCCTCCGA | |
| EsxUT-3 | CTTAATTAACGGCTGGAGCTTGGGCACGC | |
| NsiI-Dn-esxT | TGCATGCATGCAGTGCCTTGATCAACTGCTGGT | |
| Pac-SD-EsxU_1 | GCTTAATTAACAGAAAGGAGGTTAATAGTGAGCACACCGAACACGCTGA | pML4328 |
| EsxT_HindIII | CGAAGCTTCCTAGCGTGCCCAAGCTCCAGCCGCCCGC | |
| NheI-HA-EsxU-F | CTAGCTAGCAGGAGGAATTCGCCAATGTACCCGTACGACGTGCCGGACTACG CCGTGAGCACACCGAACACGCT | pML4850 |
| XmaI-EsxU-R | CTAGCCCGGGCTGATCCTCGGTCTATAGGTCGC | |
| XmaI-EsxT-F | CTAGCCCGGGGAGGATCAGCCTCGACTATGAACGCAGA | |
| HindIII-EsxT-8xHis-R | CATAAGCTTCTAGTGATGGTGATGGTGATGGTGATGGCGTGCCCAAGCTCCA GCC | |
| NdeI_Myc_esxF_F | GAACTACATATGGAACAAAAACTCATCTCAGAAGAGGATCTGATGGGTGCCGA AC | pML5056 |
| XmaI-EsxF-R | CTAGCCCGGGTCAGCCACCGCCCACCTCACGA | |
| XmaI-EsxE-F | CTAGCCCGGGAGGAGGAATTCGCCAGTGGATCCGACCGTGTTGG | |
| HindIII-EsxE-FLAG-R | CATAAGCTTTCATTTATCATCATCATCTTTATAATCCGACCACATACCCAAA TTCGT | |
| NdeI-His8-EsxF-F | GAACTACATATGCATCACCATCACCATCACCATCAC | pML5040 |
| HindIII-EsxT-Myc-R | CATAAGCTTCTACAGATCCTCTTCTGAGATGAGTTTTTGTTCGCGTGCCCAA GCTCCAGC | pML4845 |
| PacI-SD-Myc-F | GTCATTAATTAACAGAAAGGAGGTTAATAATGGAGCAGAAGCTGATCTCGGA AGAGGAC | pML4862 |
| Myc- S(G)4-EsxU-F | ATGGAGCAGAAGCTGATCTCGGAAGAGGACCTGTCCGGCGGTGGCGGTGTGA GCACACCG | |
| S(G)4-EsxU-F | TCCGGCGGTGGCGGTGTGAGCACACCGAACACGCT | |
| HindIII-EsxT-R | CATAAGCTTGCGTGCCCAAGCTCCAGCC | |
| PacI-SD-His8-F | GTCATTAATTAACAGAAAGGAGGTTAATAATGCATCACCATCACCATCACCA TCAC | pML5084 |
| His8-S(G)4-EsxF-F | ATGCATCACCATCACCATCACCATCACTCCGGCGGTGGCGGTATGGGTGCCG ACGAC | |
| HindIII-EsxE-R1 | CATAAGCTTTCACGACCACATACCCAAATTCGTGGCCATCGC | |
| PacI-SD-HA-F | GTCATTAATTAACAGAAAGGAGGTTAATAATGTACCCGTACGACGTGCCGGA CTACGCC | pML4863 |
| HA-linker_i_EsxB_FP | GTACGACGTGCCGGACTACGCCTCCGGCGGTGGCGGTATGGCAGAGATGAAG ACC | |
| EsxA-HindIII-RP | GCTAGCAAGCTTATGCGAACATCCCAGTG | |

**Table S3. Oligonucleotides used in this study**
The underlined sequences denote the restriction sites.

| | | |
|---|---|---|
| Pac-SD-EsxBF | GTCATCTTAATTAACAGAAAGGAGGTTAATAATGGCAGAGATGAAG | pML4864 |
| HindIII-HA-linker-EsxARP | GTAAAGCTTAGGCGTAGTCCGGCACGTCGTACGGGTAACCGCCACCGCCGGATGCGAACATCCCAGT | |
| HA- S(G)4-EsxD-F | ATGTACCCGTACGACGTGCCGGACTACGCCTCCGGCGGTGGCGGTGTGGCAGACACAATT | pML4854 |
| HindIII-EsxC-R-1 | CATAAGCTTAGAACAAGCCCGCGATGGCCTGG | |
| PacI-SD-EsxD-F | GTCATTAATTAACAGAAAGGAGGTTAATAGTGGCAGACACAATTCAGGTAACAC | pML4855 |
| S(G)4-EsxC-R | ACCGCCACCGCCGGAGAACAAGCCCGCGATGGCCT | |
| HindIII-HA-S(G)4-R | CATAAGCTTAGGCGTAGTCCGGCACGTCGTACGGGTAACCGCCACCGCCGGA | |
| HA- S(G)4-EsxG-F | ATGTACCCGTACGACGTGCCGGACTACGCCTCCGGCGGTGGCGGTATGAGCCTTTTGGAT | pML4860 |
| EsxH-HindIII-R | CATAAGCTTCTAGCCGCCCCATTTGGC | |
| PacI-SD-EsxG-F | GTCATTAATTAACAGAAAGGAGGTTAATAATGAGCCTTTTGGATGCTCATATCCCAC | pML4861 |
| S(G)4-EsxH-R | ACCGCCACCGCCGGAGCCGCCCCATTTGGC | |
| EsxDC-1 | TGACTAGTCCGTGGTATCGACCGCGATC | pML4805 |
| SwaI-esxDC-2-in-frame | ATTTAAATTGTTACCTGAATTGTGTCTGCCAC | |
| PacI-esxDC-3-in-frame | CTTAATTAAATCGGAACCGACCAGGCCAT | |
| EsxDC-4 | TGCATGCATGGTGTCGGATACCGAATCCC | |
| PacI-SD-EsxD-F | GTCATTAATTAACAGAAAGGAGGTTAATAGTGGCAGACACAATTCAGGTAACAC | pML4852 |
| HindIII-EsxC-R | CATAAGCTTAGAACAAGCCCGCGATGG | |
| SpeI-Up-eccCa1 | TGACTAGTCACAGTCACACGAAGCGGTGCTGGT | pML4803 |
| SwaI-Up-eccCa1 | ATTTAAATGCAGTGCGGCATCTTTCGACAGCA | |
| PacI-Dn-eccCb1 | CTTAATTAAGCCCAAGACCAGCTCCACCGTG | |
| NsiI-Dn-eccCb1 | TGCATGCATGCAAACAACGACGTCACCTGCTGCA | |

**Table S3. Oligonucleotides used in this study (continued)**
The underlined sequences denote the restriction sites.

| Antibody | Source | Identifier |
|---|---|---|
| Rabbit polyclonal anti-TNT | ProMab Biotechnologies, negatively purified antiserum was used | This study |
| Rabbit polyclonal anti-EsxU | GenScript | This study |
| Rabbit polyclonal anti-GlpX | UAB Hybridoma core | (Speer et al., 2015) |
| Mouse monoclonal anti-MctB | UAB Hybridoma core | (Speer et al., 2015) |
| Rabbit polyclonal anti-EsxT | ProMab Biotechnologies | This study |
| Rabbit polyclonal anti-EsxF | GenScript | (Tak et al., 2021) |
| Mouse monoclonal anti His-Tag | ABclonal | AE003 |
| Mouse monoclonal anti HA-Tag | Invitrogen | PI26183 |
| Mouse monoclonal anti Myc-Tag | Invitrogen | MA121316 |
| Rabbit polyclonal anti-FLAG | Millipore | F7425 |
| Mouse monoclonal anti-Galectin-3 (clone A3A12) | Abcam | ab2785 |
| Rabbit polyclonal anti-*Mycobacterium tuberculosis* | Abcam | ab905 |
| Rabbit polyclonal anti-ESAT6 | Abcam | ab45073 |
| Rabbit polyclonal anti-Antigen-85 complex | BEI Resources | NR-13800 |
| Donkey anti-rabbit IgG (H+L)-Alexa Fluor-594 secondary | Invitrogen | A-21207 |
| Goat anti-mouse IgG (H+L)-Alexa Fluor-594 secondary | Invitrogen | A-11005 |
| Donkey anti-rabbit IgG (H+L)-Alexa Fluor-488 | Invitrogen | A-21206 |
| IRDye 680RD Donkey anti-rabbit IgG secondary | LI-COR | 926-68073 |
| 6x-His Tag Monoclonal Antibody (HIS.H8), Alexa Fluor 555 | Invitrogen | MA121315A55 |
| c-Myc Monoclonal Antibody (9E10), Alexa Fluor 647 | Invitrogen | MA1980A647 |

**Table S4. Antibodies used in this study**

**a**

| **Canonical heterodimers** | **ipTM score** |
|---|---|
| EsxUT | 0.82 |
| EsxBA | 0.77 |
| EsxDC | 0.76 |
| EsxFE | 0.62 |
| EsxGH | 0.56 |

**b**

| **Non-Canonical heterodimers** | **ipTM score** |
|---|---|
| EsxFH | 0.65 |
| EsxUE | 0.60 |
| EsxDE | 0.50 |
| EsxBE | 0.47 |
| EsxGT | 0.44 |
| EsxDA | 0.43 |
| EsxDT | 0.42 |
| EsxFA | 0.42 |
| EsxGC | 0.41 |
| EsxUA | 0.40 |
| EsxBC | 0.36 |
| EsxGA | 0.36 |
| EsxUH | 0.34 |
| EsxBH | 0.33 |
| EsxBT | 0.33 |
| EsxUC | 0.30 |
| EsxGE | 0.26 |
| EsxFT | 0.25 |
| EsxFC | 0.22 |
| EsxDH | 0.18 |

**Table S5:** The ipTM scores of (**a**) Canonical and (**b**) Non-Canonical heterodimers of small Esx proteins indicate the accuracy of each predicted protein complex. The ipTM score of EsxUE is highlighted in yellow.

#### Supplementary References

Chang, A.C., and Cohen, S.N. (1978). Construction and characterization of amplifiable multicopy DNA cloning vehicles derived from the P15A cryptic miniplasmid. J Bacteriol *134*, 1141-1156.

Guilhot, C., Gicquel, B., and Martin, C. (1992). Temperature-sensitive mutants of the *Mycobacterium* plasmid pAL5000. FEMS Microbiol Lett *77*, 181-186.

Labidi, A., Mardis, E., Roe, B.A., and Wallace, R.J., Jr. (1992). Cloning and DNA sequence of the *Mycobacterium fortuitum var fortuitum* plasmid pAL5000. Plasmid *27*, 130-140.

Malagon, F. (2013). RNase III is required for localization to the nucleoid of the 5' pre-rRNA leader and for optimal induction of rRNA synthesis in E. coli. Rna *19*, 1200-1207.

Mitra, A., Speer, A., Lin, K., Ehrt, S., and Niederweis, M. (2017). PPE surface proteins are required for heme utilization by *Mycobacterium tuberculosis*. MBio *8*, e01720.

Ofer, N., Wishkautzan, M., Meijler, M., Wang, Y., Speer, A., Niederweis, M., and Gur, E. (2012). Ectoine biosynthesis in *Mycobacterium smegmatis*. Appl Environ Microbiol *78*, 7483-7486.

Pajuelo, D., Tak, U., Zhang, L., Danilchanka, O., Tischler, A.D., and Niederweis, M. (2021). Toxin secretion and trafficking by *Mycobacterium tuberculosis*. Nat Commun *12*, 6592.

Pym, A.S., Brodin, P., Brosch, R., Huerre, M., and Cole, S.T. (2002). Loss of RD1 contributed to the attenuation of the live tuberculosis vaccines *Mycobacterium bovis* BCG and *Mycobacterium microti*. Mol Microbiol *46*, 709-717.

Sampson, S.L., Dascher, C.C., Sambandamurthy, V.K., Russell, R.G., Jacobs, W.R., Jr., Bloom, B.R., and Hondalus, M.K. (2004). Protection elicited by a double leucine and pantothenate auxotroph of *Mycobacterium tuberculosis* in guinea pigs. Infect Immun *72*, 3031-3037.

Speer, A., Sun, J., Danilchanka, O., Meikle, V., Rowland, J.L., Walter, K., Buck, B.R., Pavlenok, M., Holscher, C., Ehrt, S.*, et al.* (2015). Surface hydrolysis of sphingomyelin by the outer membrane protein Rv0888 supports replication of *Mycobacterium tuberculosis* in macrophages. Mol Microbiol *97*, 881-897.

Tak, U., Dokland, T., and Niederweis, M. (2021). Pore-forming Esx proteins mediate toxin secretion by *Mycobacterium tuberculosis*. Nat Commun *12*, 394.
